# Unraveling the metabolic landscape of alkaptonuria through a human-relevant *in vitro* liver disease model

**DOI:** 10.64898/2026.09.27.754319

**Authors:** Matthias Rombaut, Gigly Del’haye, Sien Lequeue, Brendan P Norman, Liesbeth Desmet, Nina Salvi, Anja Heymans, Jessie Neuckermans, Juliette H Hughes, Gamze Ates, Ann Massie, Tamara Vanhaecke, George Bou-Gharios, Joery De Kock

## Abstract

Alkaptonuria (AKU) is a rare inherited metabolic disorder of tyrosine catabolism caused by a deficient homogentisate 1,2-dioxygenase (HGD) enzyme. This results in the accumulation of homogentisic acid (HGA), driving a progressive multisystem pathology characterized by debilitating early-onset osteoarthritis due to connective tissue degeneration. While previous *in vitro* studies have primarily relied on exogenous HGA exposure in osteoarticular cell models, the direct metabolic consequences of endogenous HGD deficiency within its native hepatic context remain poorly understood. Here, we established the first human-relevant HGD knockout hepatic *in vitro* model using a universal in-house-developed homology-directed repair approach. Integrative multi-omic analysis revealed that HGD deficiency induces widespread metabolic rewiring extending beyond disrupted tyrosine catabolism. HGD-deficient hepatocytes exhibited elevated oxidative stress accompanied by impaired mitochondrial respiration and a pseudohypoxic metabolic adaptation toward increased glycolytic dependency. Despite this glycolytic shift, the cells displayed reduced anabolic and translational activity alongside attenuated proliferation, consistent with a chronic stress-adaptive survival state rather than a proliferative metabolic phenotype. This study provides systems-level insights into the pathophysiology of AKU and establishes a versatile platform for mechanistic and therapeutic investigation.

## 2. Introduction

Alkaptonuria (AKU) (OMIM #203500) is a rare congenital liver disorder that is characterized by a deficient homogentisate 1,2-dioxygenase (HGD) enzyme. This deficiency results in the build-up of the upstream oxidation-sensitive metabolite homogentisic acid (HGA). Although the majority of circulating HGA is eliminated through urinary excretion, the remaining fraction deposits in the extracellular matrix of connective tissues as an ochronotic pigment, a process known as ochronosis^1^. Based on the oxidation properties of hydroquinone-like structures, such as HGA, the formation of benzoquinone acetic acid (BQA) has been proposed^2^. However, direct experimental evidence supporting the formation of BQA from HGA is lacking. Recent molecular spectroscopy studies showed that ochronotic pigment formation is driven by a radical-mediated oxidative coupling via the hydroxyl groups on the aromatic ring of HGA^3^. Yet the precise composition and structure of the ochronotic pigment remain up for debate, presenting a significant gap in the understanding of AKU pathophysiology^4^. Its deposition initiates a plethora of systemic manifestations, including inflammation, oxidative stress, and collagen disruption, which collectively contribute to the hallmark features of AKU, such as skin discoloration, tendon rupture and calcification, aortic stenosis, and a debilitating, early-onset form of osteoarthritis^5,6^.

*In vitro* cell models of the osteoarticular compartment showed that exogenous HGA supplementation drives oxidative stress-mediated inflammation^7,8^, proteome remodelling^9^, and structural deterioration^10^. Disruption of redox homeostasis is evidenced by elevated hydrogen peroxide levels^11^, increased lipid peroxidation^8,12,13^, accumulation of carbonylated proteins^12,13^, and decrease of free thiols^13^, alongside impairment of key antioxidant systems, including superoxide dismutase, thioredoxin, and glutathione peroxidase^11,13^. The redox imbalance is further exacerbated by increased NADPH oxidase activity, mitochondrial dysfunction, and impaired Nrf2-DNA binding^11^. Concomitantly, HGA induces inflammatory and amyloidogenic responses^7,8^, including increased oxide production and upregulation of inducible nitric oxide synthase and cyclooxygenase-2^8^. These changes are accompanied by growth inhibition^14,15^ and morphological alterations such as cytoskeletal reorganization and extracellular matrix remodelling^10^. Although informative, these models rely on exogenous HGA supplementation and therefore do not reproduce the endogenous metabolic consequences of HGD deficiency.

Urinary metabolic profiling of an AKU mouse model showed that HGA is metabolized predominantly via phase II biotransformation, where phase I detoxification is relatively minor^16^. This was also confirmed in urine of AKU patients. Interestingly, HGD deficiency induced changes to peripheral neurotransmitter metabolism, despite seeing no changes in the other upstream metabolites like tyrosine, hydroxyphenylpyruvic acid or hydroxyphenyllactic acid^16^. In addition, a shift in purine metabolism was described, hypothesized as a protective mechanism against HGA-induced oxidative stress^16^. Similarly, the serine, glycine, and one-carbon metabolism is affected, known for its role in maintenance of redox homeostasis^17^. Perturbations in metabolites linked to the tricarboxylic acid (TCA) cycle indicate a malfunction and/or a change in energy metabolism^16,17^. Recently, it was shown that tryptophan metabolism, glutamine/glutamate metabolism, histidine metabolism, citrate cycle, vitamin B6 metabolism, arginine and proline metabolism, and nicotinate metabolism are also affected by HGD deficiency^18^. These findings demonstrate that AKU is not solely a defect of tyrosine catabolism but a multisystem metabolic disorder.

Despite AKU originating in the liver, the cellular and molecular consequences of HGD deficiency at the hepatic level remain poorly understood. To address this knowledge gap, we report the first human-relevant *in vitro* liver disease model of AKU. Through integration of transcriptomic and high-resolution metabolomic profiling, we define the metabolic signatures of HGD deficiency in a liver-derived HepG2 model. This versatile platform enables the identification of (compensatory) pathway perturbations and supports the discovery and evaluation of potential therapeutic targets.

## 3. Material & Methods

### 3.1. Generation of HGD KO HepG2 clones

HepG2 cell lines and clones in this study were verified to be negative for mycoplasma contamination (Lonza MycoAlert). HepG2 cells (obtained from ATCC, HB-8065) were routinely cultivated in DMEM, high glucose, pyruvate (Gibco) supplemented with 10% (v/v) FBS, and split once or twice a week at 1:4-1:5 depending on confluence. In this study, two elongation factor 1α (EF1A) promoter-driven selection cassette plasmids were developed carrying a different antibiotic resistance and fluorescent marker followed by a stop codon; which can be made gene specific by a simple PCR to add homology arms (30-40bp)^19^. These can be noted as follows: EF1A-PuroR-2A-emGFP-STOP & EF1A-BsdR-2A-mApple-STOP (See Suppl. Doc. 1). The promotor and both selection markers are floxed and can thus be excised via Cre-lox recombination. The gene-specific selection cassettes were then co-transfected with a ribonucleoprotein CRISPR gene-editing complex via lipofectamine CRISPRMAX-mediated delivery to generate a gene knockout (KO) cell line. The Invitrogen TrueDesign Genome Editor online tool was used to design the gRNA and Invitrogen TrueCut Cas9 Protein v2 to generate the double strand break (DSB). The CRISPR-Cas9 genome editing protocol was followed according to the Invitrogen TrueGuide Synthetic gRNA manual with minor modifications. Briefly, for a 24-well format, 1250 ng Cas9 protein, 7.5 pmol gRNA, 2.5 µL Lipofectamine Cas9 Plus Reagent, and 500 ng of both selection cassettes was added to OptiMEM to a final volume of 23.5 µL. In another tube, 1.5 µL Lipofectamine CRISPRMAX Reagent was added to 25 µL of OptiMEM, vortexed and incubated at room temperature for 1 min. The former tube was added to the latter, vortexed and incubated at room temperature for 15 min. 50 000 HepG2 cells were added to this mixture, HepG2 medium supplemented with 1 µM of Alt-R HDR Enhancer V2 (IDT) was added to increase homology-directed repair (HDR) efficiency, and the cells were plated. After 24 h, the CRISPR-Cas medium was discarded, cells were rinsed and cultivated in basal HepG2 medium. After 72 h, initial HDR efficiency can be verified via fluorescence microscopy and dual antibiotic selection was started by supplementing basal HepG2 medium with 2 µg/mL puromycin and 20 µg/mL blasticidin. Single cell clones were obtained after FACS-sorting (FACSMelody, BD Biosciences) of double emGFP and mApple positive cells. The excision of the exogenous promotor and selection markers of the selection cassettes was done via lipofectamine-mediated transfection with JetMESSENGER (Polyplus) of cellulose-purified methylated Cre recombinase mRNA (Laboratory for Molecular and Cellular Therapy, VUB).

### 3.2. Validation KO state of the HGD KO HepG2 clones

Genomic DNA (gDNA) was isolated from the wild type (wt) HepG2 cell line and the HGD KO HepG2 clones using DirectPCR Lysis Reagent (Viagen Biotech) according to the manufacturer’s instructions with modifications. Cells were harvested with a mixture of Lysis Reagent supplemented with 0.4 mg/mL Proteinase K. DNA extraction was finalized by 15 min incubation at 68°C followed by 10 min at 95°C. A gene amplicon PCR was developed to verify the correct insertion of the selection cassettes in the different HGD KO HepG2 clones, and thus the KO state of the clones. Briefly, Forward (Fw) and Reverse (Rv) primers were designed to amplify an amplicon of the HGD gene, and a junction Rv (Jc) primer that is universal for both selection cassettes. A PCR mixture was made consisting of 2 µL cell lysis, 2 µL of 10 µM primer mix, 10 µL of 360 GC Enhancer (Thermo Fisher Scientific), 25 µL of 2X Phusion Flash High-Fidelity PCR Master Mix (Thermo Fisher Scientific), and 11 µL of UltraPure DNase/RNase-Free Distilled Water (Thermo Fisher Scientific). PCR was run at following conditions: 10 sec at 98°C (1 cycle), 1 sec at 98°C, 5 sec at 55°C, and 1 min at 72°C (35 cycles), and 2 min at 72°C (1 cycle). PCR products were visualized on a 2% (m/v) agarose gel stained with GelRed (Biotium) using a ChemiDoc MP imaging system (Bio-Rad).

Protein samples were taken from the wt HepG2 cell line and the HGD KO HepG2 clones and analyzed with Western Blot. Briefly, cells were harvested in RIPA buffer supplemented with 1% (v/v) Halt Protease Inhibitor Cocktail (100X) (Thermo Fisher) and 1% (v/v) 0.5 M EDTA (Thermo Fisher Scientific). The cell mixture was kept on ice for 20 min, while being vortexed shortly every 5 min. After centrifugation (13 000g, 20 min, 4°C), the protein concentration of the supernatant was determined with the Pierce BCA Protein Assay Kit (Thermo Fisher) according to the manufacturer’s protocol. After separation using sodium dodecyl sulfate (SDS) polyacrylamide gel electrophoresis (PAGE) on a 12% Mini-Protean TGX Stain-free gel (Bio-Rad), the proteins were immediately transferred onto a nitrocellulose Trans-Blot membrane (Bio-Rad). After blocking for 1 h with a solution containing 5% (m/v) milk powder in Tris-buffered saline solution with 0.1% (v/v) Tween-20, the membranes were incubated overnight at 4°C with 2 µg/mL primary polyclonal anti-HGD antibody (PA5-79360) produced in rabbits (Sigma-Aldrich) diluted in blocking buffer. After extensive washing with Tris-buffered saline solution containing 0.1% (v/v) Tween-20, the membrane was incubated at room temperature for 1 h with 0.25 µg/mL polyclonal secondary goat anti-rabbit antibody (P0448, Dako) diluted in blocking buffer, and washed with Tris-buffered saline solution containing 0.1% (v/v) Tween-20 to remove excess antibody. Finally, the HGD protein bands were detected using the Pierce enhanced chemiluminescence substrate Western Blotting substrate kit (Thermo Fisher Scientific) and ChemiDoc MP imaging system (Bio-Rad).

### 3.3. Whole genome sequencing

gDNA was extracted from wt HepG2 cells and HGD KO HepG2 clone 4 using the MagMAX™ DNA Multi-Sample Kit (Applied Biosystems) according to the manufacturer’s protocol. Libraries were prepared using the NEBNext Ultra II FS PCR FREE kit with the unique dual indexed adapters supplied in the MGIEasy Fast PCR-FREE FS V2.0 kit. The Qubit 2.0, using Promega’s Quantifluor ONE kit, and AATI Fragment Analyzer, using Agilent’s DNF-474 High Sensitivity NGS Fragment Analysis Kit, were used to quantify and qualify the library, respectively. A total of 800-1000 fmol library is denatured, circularized and digested to circular ssDNA using the MGIEasy Dual-barcode Circularization kit (MGI Tech). Subsequently, DNA Nanoballs (DNBs) were generated according to the protocol of DNBSEQ DNB Rapid Make Reagent Kit (MGI Tech).

Next, the flowcell was loaded with the DNBs using the MGI DNB Loader device and sequenced on the MGI DNBSEQ-T7 using the DNBSEQ-T7RS High Throughput Sequencing Set V3.0 kit and the High-Throughput Pair-End Sequencing Primer Kit (MGI Tech), generating sufficient paired-end 150bp reads to attain a 30x coverage.

Quality of raw sequencing reads was assessed using FastQC v0.12.1 and trimmed with Trimmomatic v0.39. Next, reads were aligned to the Homo_sapiens.GRCh38 reference genome, for which an index was generated using the bwa-mem2 index command, followed by read alignment with bwa-mem2 v2.2.1. SAM files were converted to BAM format, sorted, and indexed using SAMtools v1.22.1. Duplicate reads were marked using GATK v4.5.0.0, and reference genome dictionary and FASTA index files were generated using GATK CreateSequenceDictionary and SAMtools faidx, respectively. Variant calling was performed using GATK HaplotypeCaller and variants between wt and HGD knockout sample were compared using BCFtools v1.18. Variant effects on genes and predicted protein products were annotated using snpEff v5.2c. Finally, Integrative Genomics Viewer v2.19.1 was used to visually inspect and confirm the integration of the selection cassettes within exon 3 of the HGD gene.

### 3.4. qPCR

RNA was isolated with an RNA mini kit (Qiagen) according to the manual. Briefly, cells were lysed with RNA lysis buffer containing 1% v/v β-mercaptoethanol (Sigma-Aldrich), mixed with ethanol and bound to a column. After extensively washing, total RNA was eluted and purified using the GenElute Mammalian Total RNA Purification Miniprep Kit (Sigma-Aldrich) and quantified by a Nanodrop spectrophotometer (Thermo Fisher Scientific). Complimentary DNA was generated with the iScript cDNA Synthesis Kit (Bio-Rad) and was purified using the GenElute PCR Clean-up Kit (Sigma-Aldrich). A QuantStudio 3 (Thermo Fisher Scientific) is used for reverse transcription quantitative polymerase chain reaction (RT-qPCR) using TaqMan Mastermix and primers. Normalization of data was performed against the geometric mean of housekeeping genes glucuronidase beta (GUSB), hypoxanthine phosphoribosyltransferase 1 (HPRT1), and ubiquitin C (UBC) using qBase+ software (Biogazelle).

### 3.5. Microarray profiling and analysis

RNA was isolated as described above. 100 ng RNA was *in vitro* transcribed using the Genechip Whole Transcriptome PLUS Reagent Kit according to the manufacturer’s instructions (Applied Biosystems). The amplified RNA and synthesized single-stranded cDNA were purified with magnetic beads. This was followed by RNase H treatment to hydrolyze 15 µg of ss-cDNA, after which 5.5 µg and 3.5 µg of ss-cDNA were fragmented and labeled using Fragmentation Master Mix and Labelling Master Mix, respectively. The labeled cDNA was then hybridized to Genechip Human Genome U133 array and placed in a Genechip Hybridization Oven-645 (Affymetrix), rotating at 14 g and 45°C for 16 h. After incubation, the arrays were washed on a Genechip Fluidics Station 450 (Affymetrix) and stained using the Affymetrix HWS kit. A GeneChip Scanner 3000 7G (Affymetrix) was employed to scan the microarray chips. Quality control of the chips was carried out using Affymetrix GCOS software and the datasets were corrected, summarized, and normalized via Robust Multiarray Analysis. Transcriptomic pathway analysis was conducted with Ingenuity Pathway Analysis software. Gene set enrichment was assessed based on a fold change of ≥2 and a Benjamini-Hochberg (B-H)-adjusted p-value threshold of <0.05.

### 3.6. Cell viability after HGA exposure

HepG2 cells and HGD KO HepG2 clones were seeded at 50 000 cells per well in a 96-well microtiter plate. After 24 h, the medium was replaced with fresh basal HepG2 medium supplemented with a wide concentration range of HGA (0.1, 0.25, 0.5, 1, 1.5, 2, 2.5, 3, 3.5, 4 mM). Since HGA was initially dissolved in DMSO, all wells were supplemented with DMSO to the DMSO concentration of the highest HGA concentration. Cells were incubated for 96 h and cell viability was assessed using a 3-[4,5-dimethylthiazol-2-yl]-2,5-diphenyl tetrazolium bromide (MTT) assay (Sigma-Aldrich). Briefly, cells were exposed to 0.5 mg/mL MTT solution for 90 min. After aspiration of the medium, the formazan crystals were dissolved with DMSO and absorbance was measured at 595 nm with a microplate reader (Spectramax iD3, Molecular Devices). Cells incubated with vehicle control medium were used as negative control. The 96 h concentration inducing 50% cytotoxicity (CC50) was calculated in RStudio (v2025.09.2) using the DRC package^20^ and the dose-response curve was plotted with GraphPad Prism using a four-parameter logistic nonlinear regression model (Dotmatics).

### 3.7. Oxidative stress assay

Reactive oxygen species were quantified in HepG2 cells and HGD KO HepG2 clone 4 with CellROX™ Orange Reagent (Invitrogen) according to the manufacturer’s protocol. Upon oxidation, the probe emits strong orange fluorescence that remains localized within the cell. Briefly, cells were exposed to 5µM CellROX™ Orange Reagent for 30 min at 37°C and during the last 5 min Hoechst 33342 (Invitrogen) was added. Flow cytometric analysis was performed on the Attune NxT Flow Cytometer running on Attune Cytometric Software v6.2.3 (Invitrogen). Percentage of orange fluorescent cells was used for analysis. Statistical analysis was performed using GraphPad Prism (Dotmatics).

### 3.8. Lipid peroxidation test

Lipid peroxidation was quantified in HepG2 cells and HGD KO HepG2 clone 4 with BODIPY 581/591 undecanoic acid (Invitrogen) according to the manufacturer’s protocol. Upon oxidation, the fluorescence emission peak of the probe (fatty acid analogue) shifts from red to green. Briefly, cells were exposed to 10µM probe for 30 min at 37°C and during the last 5 min Hoechst 33342 (Invitrogen) was added. Flow cytometric analysis was performed on the Attune NxT Flow Cytometer running on Attune Cytometric Software v6.2.3 (Invitrogen). The percentage of green fluorescent cells was used for analysis. Statistical analysis was performed using GraphPad Prism (Dotmatics).

### 3.9. Neutral lipids staining

Flow cytometric analysis of intracellular neutral lipids was performed as previously described^21^. Briefly, HepG2 cells and HGD KO HepG2 clone 4 were stained in suspension with 2.5µM BODIPY 493/503 Neutral Lipid Stain (Invitrogen) for 10 min. Nuclei were stained for the last 5 min with 5 µg/mL Hoechst 33342 (Invitrogen) before analysis on the Attune NxT Flow Cytometer running on Attune Cytometric Software v6.2.3 (Invitrogen). Statistical analysis was performed using GraphPad Prism (Dotmatics).

### 3.10. Real-time metabolic analysis with Seahorse Analyzer

The influence of HGD deficiency on mitochondrial function was evaluated using the MitoStress Test (Agilent Technologies) according to the manufacturer’s protocol. Briefly, HepG2 cells and HGD KO HepG2 clone 4 were plated (20 000 cells/well) in a 96-well Seahorse assay plate for 24 h under basal cultivation conditions. Before the assay, the cultivation medium was replaced with XF DMEM supplemented with 1 mM pyruvate, 2 mM glutamine, and 20 mM glucose and transferred to a CO_2_-free incubator at 37 °C for at least 1 h. The final concentrations of the reagents of the MitoStress Test were as followed: oligomycin 1 µM, FCCP (carbonyl cyanide-p-(trifluoromethoxy)phenylhydrazone) 1 µM, and Rot/AA (rotenone/antimycin A) 2.5 µM. After staining with 4 µM Hoechst 33342 (Thermo Fisher), imaging was performed with BioTek Cytation 1 Cell Imaging Multimode Reader (Agilent Technologies) to normalize the data. Data was analysed using Seahorse Analytics software (Agilent Technologies) with each condition consisting of 21 samples. Statistical analysis was performed using GraphPad Prism (Dotmatics).

### 3.11. Metabolic profiling

HepG2 cells and HGD KO HepG2 clone 4 are routinely cultivated in DMEM, high glucose, pyruvate (Gibco) supplemented with 10% FBS. Importantly, this basal cultivation medium already contains 360 µM phenylalanine and 360 µM tyrosine. If stated, this medium was supplemented with 1 mM or 2 mM of phenylalanine, tyrosine or 4-hydroxyphenylpyruvic acid (4-HPP) (Sigma). Additionally, a basal cultivation medium was made deprived from phenylalanine and tyrosine. After 72 h of exposure, cell supernatant was collected and centrifuged for 5 min at 10 000g.

Analysis of cell supernatant was performed on an Acquity UPLC I-Class coupled to a Vion IMS QTOF detection system from Waters Corporation, equipped with a Z-spray electrospray ion source.

Reversed-phase liquid chromatography was performed on Waters Acquity Premier High Strength Silica T3 column (1.8 µm, 2.1×100 mm) with VanGuard FIT maintained at 40°C and a flow rate of 0.4 mL/min. Mobile phase composition was (A) Milli-Q water (MQ) and (B) methanol (MeOH; Thermo Fisher Scientific), both with 5 mM ammonium formate (AMF; Sigma) and 0.1% (v/v) formic acid (FA; Honeywell). The elution gradient began at 100% of mobile phase A for 1 min and decreased linearly over 11 min until 100% of mobile phase B was attained. This was maintained for 2 min after which the column was re-equilibrated at the initial conditions for 2 min. Protein precipitation on cell supernatant was carried out by adding 1:3 ratio of ice cold MeOH. After 10 min of incubation at 4°C, samples were centrifuged at 10 000g for 15 min at 4°C. The supernatant was subsequently dried under a stream of nitrogen and reconstituted in MQ supplemented with 0.1% (v/v) FA.

The Hydrophilic Interaction Liquid Chromatography (HILIC) separation was performed in a Waters Acquity Premier Ethylene-Bridged Hybrid Amide column (1.7 µm, 2.1 x 100 mm) with VanGuard FIT maintained at 40°C and a flow rate of 0.5 mL/min. Mobile phase A consisted of 5 mM AMF in 95:5 (v/v) ACN/MQ with 0.1% (v/v) FA, while mobile phase B consisted of 5 mM AMF in 60:40 (v/v) MQ/ACN with 0.1% (v/v) FA. The elution gradient began at 100% of mobile phase A and decreased linearly over 6 min until 60% of mobile phase B was attained. Over 1 min it was further linear increased to 100% B, which was held for 2 min, followed by re-equilibration at initial conditions for 10 min. To precipitate the proteins in the cell supernatant, 50 µL of centrifuged supernatant was added to 150 µL cold precipitation mixture consisting of 130/10/10 (v/v/v) ACN/MeOH/MQ. After 10 min of incubation at 4°C, samples were centrifuged at 10 000g for 15 min at 4°C and the supernatant was ready for analysis.

A pooled quality control sample was prepared and injected ten times at the beginning of the analytical run and repeated every tenth sample to monitor system stability. Experimental sample order was randomized prior to injection. Sample injection volume was 6 µL and the autosampler compartment was kept at 4°C. The sample wash solvent consisted of Isopropanol/MeOH/ACN/MQ (25:25:25:25 (v/v/v/v); Isopropanol, Thermo Fisher Scientific) with 0.2% (v/v) FA, while purge and seal wash solvents consisted of MeOH/MQ (20:80 (v/v)). MS^E^ data-independent acquisition was performed in negative ionisation polarity, as it showed a higher sensitivity than positive ionisation mode for our in-house library. The scan settings were set to an m/z range of 50 to 1000, with scan time set to 0.2 seconds. The collision energy was set at 6eV for low energy and ranging from 28 to 56eV for high-energy fragmentation. Mass accuracy was maintained using continuous infusion of Leucine enkephalin (150 pg/µL in 50:50 MQ/ACN with 0.1% (v/v) FA) as lock mass (m/z 554.2615 [M−H]⁻).

Retention time alignment, peak picking, noise reduction, and metabolite annotation was performed with Progenesis QI (version 2.3). The following adducts were considered during data processing: [M−H]⁻, [M+FA−H]⁻, and [M−H₂O−H]⁻. Metabolite abundance were normalised to all compounds and the normalised abundance was used to perform statistical analysis using GraphPad Prism (Dotmatics). A false discovery rate (FDR)-adjusted p-value (q-value) < 0.05 was set to annotate differential metabolites.

When available, accurate mass and retention time (AMRT) was used to identify metabolites with an in-house library of 54 and 57 reference standards for RP and HILIC analysis, respectively (See Suppl. Doc 2). Identification required the metabolite to have a retention time tolerance within ±0.3 min and a mass accuracy within ±10 ppm. Other annotated metabolites were identified based on an in-house literature-based database of metabolites related to untreated AKU mice/patients solely on AM. These putative metabolites were also assigned using a mass accuracy threshold of ±10 ppm.

## 4. Results

### 4.1. Generation of a homozygous HGD KO HepG2 cell line

After lipofectamine-mediated delivery of the ribonucleoprotein CRISPR gene-editing complex with the selection cassettes (See Fig. 1A), an initial knock-in efficiency of the emGFP-based selection cassette of 2.15% was obtained (See Suppl. Fig. 1A). Subsequent dual antibiotic selection resulted in an enrichment to a mixed population of 0.46% emGFP^-^/mApple^-^, 6.78% emGFP^+^/mApple^-^, 6.13% emGFP^-^/mApple^+^, and 86.62% emGFP^+^/mApple^+^ (See Suppl. Fig. 1A). A bulk cell line of emGFP^+^/mApple^+^ cells was established and single cell cloning of emGFP^+^/mApple^+^ was performed (See Fig. 1B), resulting in the generation of sixteen different HGD KO HepG2 clones. The CRE mRNA-induced excision of the selection cassette resulted in a mixed population of 32.12% emGFP^+^/mApple^+^, 6.46% emGFP^-^/mApple^+^, 2.31% emGFP^+^/mApple^-^, and 46.64% emGFP^-^/mApple^-^(See Suppl. Fig. 1A).

**Fig. 1:**
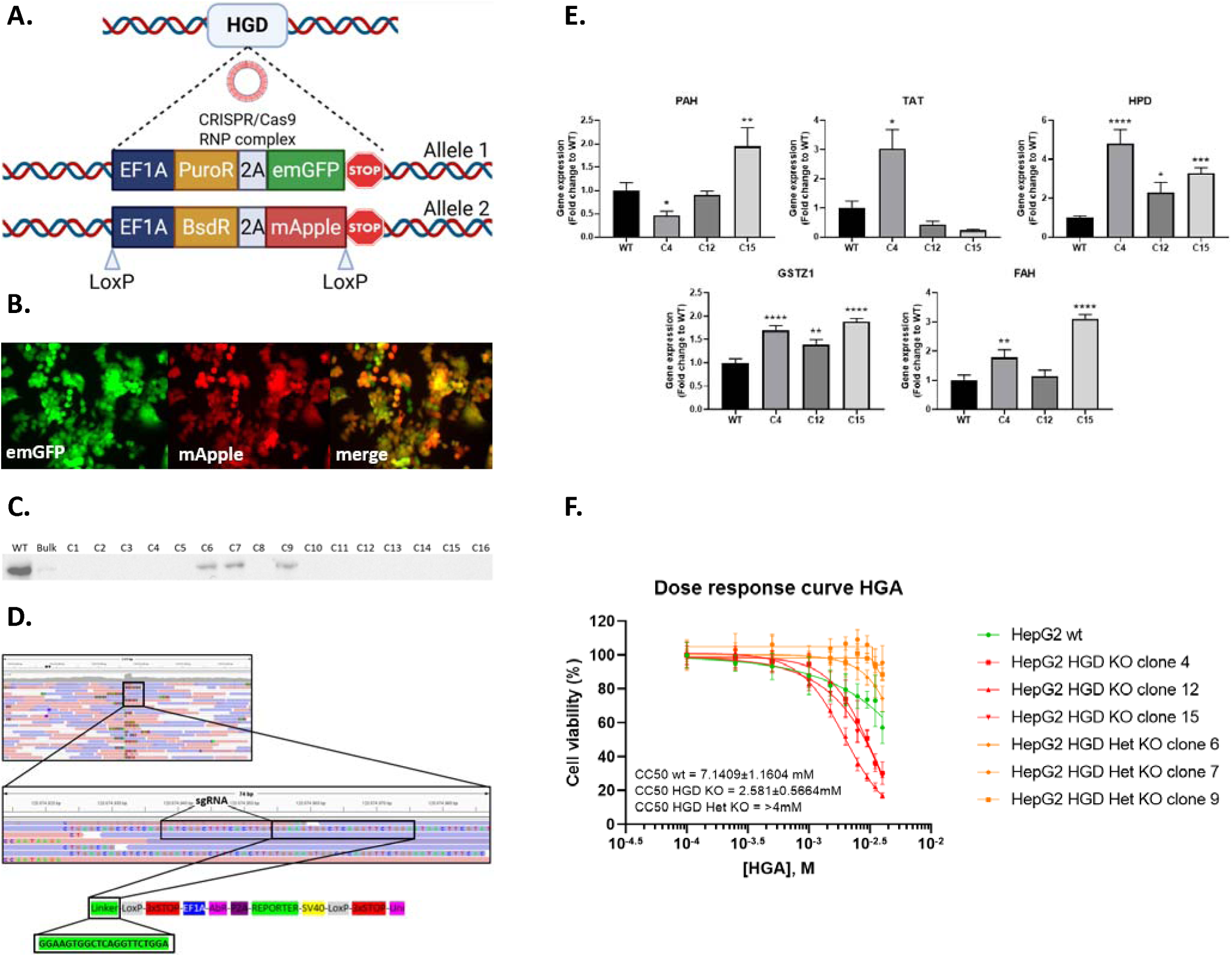
Generation and characterization of HGD-deficient HepG2 cells. **A:** Visual representation of the gene KO method with both selection cassettes integrated in the HGD gene (image made with Biorender). **B:** Fluorescent images of emGFP-and mApple-postive HepG2 cells after dual antibiotic selection (200x magnification). **C:** Representative cropped western blot for HGD in HepG2 cells: Wild-type, bulk, and sixteen HGD KO clones are consecutively blotted. **D:** Whole Genome Sequencing: read alignment shows the correct integration of HDR constructs in exon 3 of HGD, next to the PAM sequence of sgRNA. **E:** Relative PAT, TAT, HPD, GSTZ1, and FAH mRNA expression in HepG2 wt and HGD KO clone 4, 12, and 15 measured by RT-qPCR and normalized to housekeeping genes GUSB, HPRT1, and UBC under basal cultivation conditions of tyrosine. One-way ANOVA Tukey’s multiple comparison test): *:p < 0.05, **: p < 0.01, ***: p < 0.001, ****: p < 0.0001 (n=3). **F:** Cell viability assay on wild-type heterozygous KO, and HGD-deficient HepG2 cells after exposure to a wide concentration range of HGA (n=8).

A gene amplicon PCR was developed to verify the correct insertion of the selection cassettes in the different HGD KO HepG2 clones, and thus the KO state of the HGD gene (See Suppl. Fig. 1B). Briefly, an amplicon of approximately 300 bp is formed when there is still unmodified HGD gene present (lane 1: Fw & Rv primer). When there is a correct integration of a selection cassette, an amplicon of approximately 500 bp is formed (lane 2: Fw & Jc primer). In lane 3, the combination of Fw, Rv and Jc primer would result in one band at approximately 300 bp if only the unmodified HGD gene is present, two bands when there is a heterozygous KO and one band at approximately 500 bp if both selection cassettes are correctly inserted in the HGD gene (=homozygous KO). As can be seen in Suppl. Fig.1B, clones 4, 12, and 15 show a homozygous KO of HGD at the gene level. Subsequent Sanger sequencing of the amplicons formed by Fw & Rv primers (lane 1) of the other clones, revealed three HGD KO HepG2 clones (C6, C7, and C9) with an intact allele, thus corresponding to a heterozygous KO of the HGD gene. Interestingly, two clones (C1 and C3) showed only a partial integration of a selection cassette and the remaining other clones had insertions or deletions of nucleotides resulting from non-homologous end joining repair mechanism (data not shown).

Western Blot analysis was performed on wt HepG2 cells, HGD KO bulk HepG2 cell line, and the sixteen HGD KO HepG2 clones (See Fig. 1C & Suppl. Fig. 1C). Interestingly, there is still functional HGD protein present in the HGD KO bulk HepG2 sample, proving that there is a chance of random integration of the selection cassettes despite homology arms specific to the HGD gene. This is further confirmed by the presence of three heterozygous HGD KO clones (C6, C7, and C9), which is in line with the gene amplicon PCR. Altogether, thirteen out of sixteen clones had a successful KO of the HGD gene, of which three with a correct integration of selection cassettes according to the gene amplicon PCR (C4, C12, and C15). For clone 4, the correct integration of both selection cassettes next to the PAM sequence of the HGD-directed gRNA was further confirmed with whole genome sequencing (See Fig. 1D). In addition, random selection cassette integrations were not found in the genome, nor chromosomal abnormalities. Although, one gene, ADGRG2, showed an acquired heterozygous mutation compared to the wt HepG2 cells.

The tyrosine catabolic pathway was analysed via qPCR on the three HGD KO HepG2 clones with correct selection cassettes integration (C4, C12, and C15) to assess the influence of HGD deficiency. HGD-deficient HepG2 cells showed an upregulation for HPD, the enzyme upstream of HGD and GSTZ1, the enzyme that comes after HGD. Interestingly, there also seems to be a balance between PAH and TAT expression (See Fig. 1E). Exposing HGD-deficient HepG2 cells to supraphysiological concentrations of tyrosine did not alter enzyme expression of the tyrosine catabolic pathway (See Suppl. Fig. 1D). In contrast, deprivation of tyrosine and phenylalanine reshaped pathway gene expression in both wt and HGD-deficient HepG2 cells. TAT expression was heavily affected, whereas HPD, GSTZ1, and FAH were mostly upregulated, eliminating the differential expression observed between wt and HGD-deficient cells under standard culture conditions (See Suppl. Fig. 1D). Homozygous and heterozygous HGD KO HepG2 clones and wt HepG2 cells were challenged to a broad concentration range of HGA. The HGD-deficient HepG2 cells showed a 2.75-fold higher susceptibility to HGA-induced cell toxicity, measured by cell viability (See Fig. 1F).

### 4.2. Transcriptomic profiling reveals a coordinated activation of glycolytic and chronic metabolic stress programs in HGD-deficient hepatic cells

Comparative transcriptomic analysis of three independent homozygous HGD KO HepG2 clones (C4, C12, and C15) *versus* wt HepG2 cells identified extensive transcriptional remodelling with 844 genes upregulated and 1037 genes downregulated. Focusing on genes exhibiting >10-fold differential expression revealed a pronounced metabolic reprogramming (See Fig. 2A and Suppl. Table 1). Notably, the glucose transporters SLC2A3 and SLC2A14 were strongly upregulated, indicating a substantial increase in glucose import. This was accompanied by induction of PFKFB3, a key regulator of glycolytic flux through fructose-2,6-bisphosphate production, consistent with a shift toward glycolysis. In parallel, upregulation of CA9, a regulator of intracellular pH, supports the presence of a glycolysis-associated acid load. CA9 also acts as a transcriptional target of hypoxia-inducible factor (HIF) signaling and its induction was associated with an increased expression of EGLN3, a key regulator of HIF signaling. Upregulation of SLC6A8 further suggests an additional response to metabolic stress to buffer cellular energy demands by enhancing creatine uptake. Induction of NDRG1, a well-established stress-responsive gene, further supports the presence of a chronic cellular stress state in the AKU model. In line with this, upregulation of RAP1A and FAM13A, both associated with small GTPase-associated signaling, LOX, and TUBB1 point to extensive remodelling of cellular signaling networks via cytoskeletal modifications and stress-responsive signaling. Surprisingly, downregulation of SLC7A11, which mediates extracellular cystine import for glutathione synthesis, combined with reduced expression of CTH, limiting intracellular cysteine production, indicates a coordinated impairment of cysteine availability. Consistently, downregulation of ALDH1L2, a mitochondrial enzyme involved in folate metabolism and redox balance, further suggests mitochondrial vulnerability. The downregulation of ASNS indicates an impaired asparagine biosynthesis with secondary consequences for glutamate metabolism. Together, these changes predict a reduced capacity to sustain antioxidant defenses and amino acid biosynthesis under stress conditions.

**Fig. 2:**
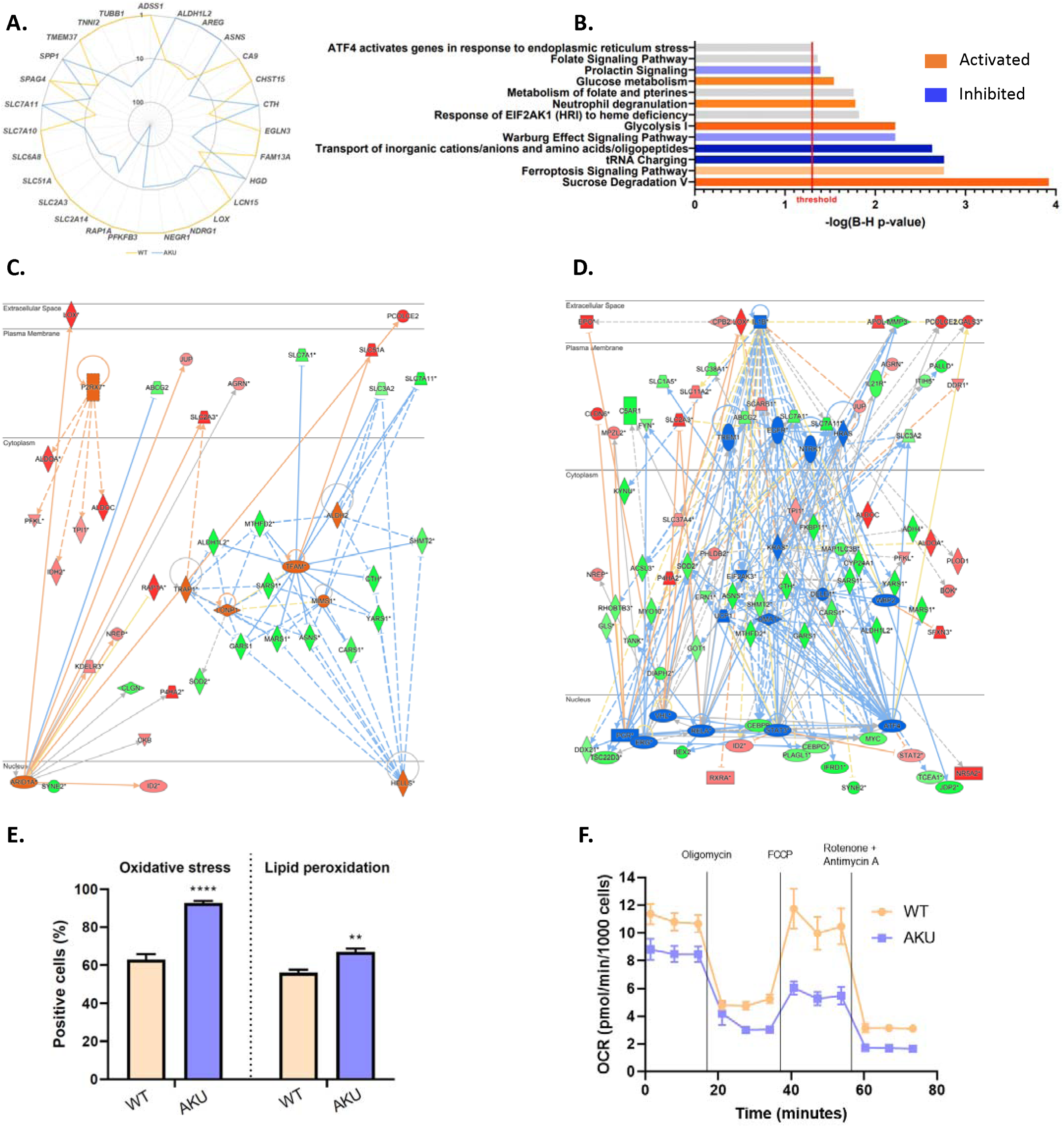
Transcriptomic and functional profiling of HGD-deficient HepG2 cells. **A:** Radar chart of transcriptome analysis of wt HepG2 cells versus HepG2 AKU model of genes with >10-fold differential expression. **B:** Canonical pathway analysis with Ingenuity Pathway Analysis of wt HepG2 cells versus HepG2 AKU model. Threshold corresponds to a q-value of <0.05, orange and blue bars show a predicted activation and inhibition of the canonical pathway, respectively. **C:** Upstream regulator analysis with Ingenuity Pathway Analysis. Visual representation of the predicted activated upstream regulators. An activation z-score of <-2 or >2 and a q-value of 0.05 were used as a threshold. **D:** Upstream regulator analysis with Ingenuity Pathway Analysis. Visual repesentation of the predicted inhibited upstream regulators. An activation z-score of <-2 or >2 and a q-value of 0.05 were used as a threshold. **E:** Flow cytometric analysis of oxidative stress and lipid peroxidation tested with CellROX™ Orange Reagent and BODIPY 581/591 undecanoic acid, respectively, in wild type HepG2 cells versus HepG2 AKU model. Student’s t-test: **: p < 0.01,****: p < 0.0001 (n=3). **F:** Seahorse mitochondrial stress test: oxygen consumption rate HepG2 cells versus HepG2 AKU model (n=21).

Ingenuity Pathway Analysis (IPA) (See Fig. 2B and Suppl. Fig. 2A) of all differentially expressed genes with a >2 absolute fold change and a q-value of <0.05 predicted a strong activation of the canonical pathways “Sucrose Degradation V”, “Glycolysis I”, and “Glucose Metabolism”, which is in line with the alteration of cellular energetics towards glycolysis. Concordantly, there is a predicted inhibition of the Warburg effect and “Prolactin Signaling”, which suggests attenuation of cell growth in the AKU model. In addition, there is a predicted inhibition of “tRNA Charging” and “Transport of inorganic cations/anions and amino acids/oligopeptides” indicating a reduction in protein synthesis and thus attenuated metabolic activity. Modulation of pathways linked to the folate cycle show widespread rewiring of one-carbon metabolism involved in nucleotide biosynthesis and redox homeostasis in response to metabolic stress. Several stress-responsive canonical pathways were significantly modulated such as the “ATF4 activates genes in response to endoplasmic reticulum stress” and the “Response of EIF2AK1 (HRI) to heme deficiency” which are both linked to the integrated stress response (ISR). Similarly, an activation of the “Neutrophil Degranulation” canonical pathway points towards oxidative stress conditions and cytoskeletal remodelling. This is further supported by a predicted activation of “Ferroptosis Signaling Pathway” that suggests a higher susceptibility to lipid peroxidation.

To define the underlying architecture regulating the observed transcriptional remodelling, IPA upstream regulator analysis was performed (See Suppl. Table 2). Among the most significantly predicted activated regulators (See Fig. 2C) are TFAM, HELLS, MIMS1, LONP1, and TRAP1, responsible for mitochondrial biogenesis, mitochondrial dynamics, regulator of mitochondrial homeostasis, mitochondrial protein quality control, and mitochondrial integrity, respectively. On the other hand, there is a predicted inhibition (See Fig. 2D) of DELE1, a critical messenger of mitochondrial stress. Interestingly, there is a coordinated predicted inhibition of EIF2AK3 (PERK) and its downstream effector ATF4, central components of the ISR program. In line with the attenuated cell growth, there is a predicted inhibition of KRAS, EGFR, STAT3, and HRAS. In addition, the predicted inhibition of RELA, STAT3, IL1B, and TREM1 shows a suppression of inflammatory signaling. The hypoxia-like transcriptional program, despite normoxic culture conditions, is influenced by a suppression of VHL.

To determine whether the transcriptional signatures showing impaired redox buffering capacity and ferroptosis susceptibility translates into functional phenotypes, intracellular oxidative stress and lipid peroxidation in HGD-deficient and wt HepG2 cells was quantified. AKU cells exhibited a marked increase in reactive oxygen species (ROS) compared to control. In parallel, lipid peroxidation levels were significantly elevated in the AKU model, indicating enhanced oxidative damage to membrane lipids (See Fig. 2D). On a side note, the levels of intracellular neutral lipids were not significantly different between both groups (See Suppl. Fig. 2B). Similarly, mitochondrial dysfunction was assessed with a Seahorse mitochondrial stress test. Strikingly, the AKU model showed a lower maximal respiration and lower basal oxygen consumption rate (OCR), resulting in limited ability to increase energy production on demand and in general less energy production via mitochondria, respectively. Additionally, they have a lower spare capacity, which makes the HGD-deficient cells more vulnerable to stress. The reduced non-mitochondrial oxygen consumption in the HepG2 AKU model indicates a decreased activity of extramitochondrial oxygen-consuming enzymes, consistent with an overall suppressed cellular metabolic state (See Fig. 2E and Suppl. Fig.2C).

### 4.3. Metabolomic profiling reveals a widespread metabolic rewiring in HGD-deficient hepatic cells

High-resolution metabolomics of conditioned media revealed extensive metabolic rewiring in the homozygous HGD KO HepG2 clone 4 compared to wt control (See Table 1). As expected from disruption of HGD, there was an accumulation of tyrosine-derived and HGA-related metabolites. Increases of BQA, tyramine-O-sulfate, γ-glutamyl-tyrosine, hydroxybenzaldehyde, tyrosol, acetyl-L-tyrosine, and p-hydroxyphenyllactic acid show an extensive rerouting of tyrosine catabolism towards alternative conjugation and metabolic pathways, potentially to mitigate HGA-induced toxicity (See Table 1).

**Table 1:** High-resolution metabolomic analysis of conditioned media of HGD-deficient HepG2 cells (AKU model) versus wild type HepG2 cells. Showing differentially abundant metabolites in Hydrophilic Interaction Liquid Chromatography (HILIC) and Reversed-phase (RP) with a q-value <0.05, identified via an in-house accurate mass and retention time (AMRT) library or an in-house literature database (AM) of metabolites related to untreated AKU mice/patients. Yellow background indicates increased metabolite abundance, whereas blue background indicates decreased metabolite abundance.

| Compound | Neutral mass (Da) | Retention time (min) | Log<sub>2</sub> FC | q-value | Identification (mass error) | Classification |
| --- | --- | --- | --- | --- | --- | --- |
| *HILIC (negative polarity)* |  |  |  |  |  |  |
| Benzoquinoneacetic acid | 166.0283 | 1.52 | ∞ | <0.0001 | AM (6.94 ppm) | HGA metabolism |
| L-arginine | 174.1118 | 5.30 | 3.94 | <0.01 | AMRT | Amino acid |
| Tyramine-O-sulfate | 217.0412 | 2.49 | 1.21 | <0.001 | AM (0.92 ppm) | Alternative tyrosine metabolism |
| Pyroglutamic acid | 129.0428 | 3.57 | 1.07 | <0.05 | AM (2.34 ppm) | Glutathione cycle |
| Adenine | 135.0547 | 2.89 | 0.61 | <0.05 | AM (2.87 ppm) | Purine metabolism |
| Indole-3-acetyl-alanine | 246.0999 | 1.82 | 0.36 | <0.05 | AM (4.53 ppm) | Indole metabolism |
| L-glutamine | 146.0694 | 4.43 | 0.28 | <0.05 | AMRT | Amino acid |
| L-phenylalanine | 165.0792 | 3.10 | -3.36 | <0.05 | AMRT | Amino acid |
| N-acetylserine | 147.0534 | 3.37 | -1.30 | <0.05 | AM (2.50 ppm) | Acetylated amino acid |
| Hydroxyisovaleric acid | 118.0632 | 1.38 | -0.89 | <0.05 | AMRT | Branched-chain amino acid |
| L-acetylcarnitine | 203.1158 | 1.84 | -0.55 | <0.05 | AM (2.42 ppm) | Mitochondrial biomarker |
| L-lysine | 146.1058 | 5.40 | -0.48 | <0.05 | AMRT | Amino acid |
| *RP (negative polarity)* |  |  |  |  |  |  |
| Benzoquinoneacetic acid | 166.0282 | 5.47 | ∞ | <0.0001 | AM (6.04 ppm) | HGA metabolism |
| Methylxanthine | 166.0478 | 1.84 | 1.40 | <0.05 | AM (7.16 ppm) | Purine metabolism |
| Malic acid | 134.0217 | 0.84 | 1.28 | <0.05 | AM (0.04 ppm) | TCA cycle |
| Ascorbic acid | 176.0321 | 0.86 | 1.18 | <0.05 | AM (2.92 ppm) | Antioxidant |
| Isocitric acid | 192.0270 | 1.10 | 1.15 | <0.01 | AM (2.80 ppm) | TCA cycle |
| γ-Glutamyl-tyrosine | 310.1162 | 3.58 | 0.95 | <0.05 | AM (2.59 ppm) | Tyrosine metabolism |
| Cholic acid | 408.2877 | 10.88 | 0.89 | <0.05 | AMRT | Bile acid |
| Hydroxybenzaldehyde | 122.0370 | 5.09 | 0.87 | <0.05 | AM (2.49 ppm) | Alternative tyrosine metabolism |
| Pyroglutamic Acid | 129.0429 | 1.48 | 0.80 | <0.01 | AMRT | Glutathione cycle |
| Hydroxybutyric acid | 104.0476 | 2.41 | 0.71 | <0.001 | AM (3.15 ppm) | Metabolic stress marker |
| Guanosine | 283.0915 | 3.01 | 0.60 | <0.05 | AMRT | Purine metabolism |
| Uric acid | 168.0283 | 1.56 | 0.55 | <0.01 | AMRT | Purine metabolism |
| N-acetyl-L-tyrosine | 223.0845 | 4.34 | 0.54 | <0.0001 | AM (2.14 ppm) | Tyrosine metabolism |
| Inosine | 268.0807 | 2.97 | 0.38 | <0.01 | AMRT | Purine metabolism |
| Tyrosol | 138.0683 | 4.23 | 0.33 | <0.05 | AM (2.62 ppm) | Catechol metabolism |
| Ketoleucine | 130.0633 | 4.34 | 0.32 | <0.05 | AM (2.24 ppm) | Branched-chain amino acid |
| L-glutamine | 146.0693 | 0.60 | 0.26 | <0.05 | AMRT | Amino acid |
| p-Hydroxyphenyllactic acid | 182.0581 | 4.23 | 0.26 | <0.05 | AMRT | Tyrosine metabolism |
| Hippuric acid | 179.0584 | 4.73 | 0.23 | <0.05 | AM (2.18 ppm) | Modulator of metabolic acidosis |
| Indole-3-acetyl-alanine | 246.1005 | 6.13 | 0.22 | <0.05 | AM (2.11 ppm) | Indole metabolism |
| Indoleacetic acid | 175.0634 | 5.86 | 0.22 | <0.01 | AM (2.81 ppm) | Indole metabolism |
| Allysine | 145.0741 | 2.81 | 0.20 | <0.05 | AM (2.66 ppm) | Amino acid derivative |
| Kynurenic acid | 189.0430 | 6.33 | 0.18 | <0.05 | AM (0.89 ppm) | Tryptophan metabolism |
| Indole-3-carboxaldehyde | 145.0531 | 4.65 | 0.13 | <0.05 | AM (1.58 ppm) | Indole metabolism |
| L-phenylalanine | 165.0790 | 3.34 | -6.67 | <0.001 | AMRT | Amino acid |
| L-methionine | 149.0512 | 1.23 | -2.40 | <0.01 | AMRT | Amino acid |
| L-glutamic acid | 147.0535 | 0.62 | -2.07 | <0.05 | AMRT | Amino acid |
| N-acetylserine | 147.0534 | 2.45 | -1.29 | <0.01 | AM (2.64 ppm) | Acetylated amino acid |
| N-acetylaspartic acid | 175.0482 | 1.14 | -1.09 | <0.01 | AMRT | Acetylated amino acid |
| Uridine | 244.0693 | 2.44 | -1.01 | <0.05 | AM (3.21 ppm) | Pyrimidine metabolism |
| Hydroxyisovaleric acid | 118.0632 | 3.99 | -0.78 | <0.01 | AMRT | Branched-chain amino acid |
| L-acetylcarnitine | 203.1158 | 6.22 | -0.57 | <0.01 | AM (2.42 ppm) | Mitochondrial fatty acid oxidation |
| O-succinyl-L-homoserine | 219.0742 | 2.49 | -0.49 | <0.01 | AM (3.14 ppm) | Amino acid derivative |
| N-acetylneuraminic acid | 309.1060 | 0.67 | -0.30 | <0.05 | AMRT | Modulator glycan function |

A central feature of the AKU metabolic landscape was the differential abundance of metabolites linked to oxidative stress and redox-buffering systems. Pyroglutamic acid and hydroxybutyric acid levels were elevated, indicating glutathione depletion and increased oxidative stress, respectively. Elevated ascorbic acid levels further supports higher demand to antioxidant mechanisms. Elevated TCA cycle intermediates (malic acid and isocitric acid) indicate mitochondrial stress. The changes in branched-amino acid derivatives (hydroxyisovaleric acid and ketoleucine) further point toward a metabolic rewiring affecting mitochondrial function (See Table 1).

Another notable feature of the AKU model is the substantial rerouting of several amino acids. Interestingly, a significant increase of L-arginine is observed, indicative of oxidative stress through formation of reactive nitrogen species via NOS system. This amino acid also serves as an intermediate in the urea cycle, a pathway that runs in parallel with the TCA cycle. Similarly, metabolites derived from tryptophan and indole metabolism were elevated, suggesting metabolic adaptation due to oxidative stress signaling. Consistent decrease in N-acetylated amino acids indicates lower levels of acetyl-CoA, which closely tracks with an altered energy demand. Mixed altered presence of purine and pyrimidine metabolites suggests a remodelling of nucleotide turnover, also reflecting altered energy balance and cellular growth dynamics (See Table 1).

Wt and HGD-deficient HepG2 cells were also challenged with increased concentrations of phenylalanine, tyrosine, and 4-HPP, or cultured under phenylalanine and tyrosine-deprived conditions to distinguish substrate-dependent metabolic alterations from HGD deficiency-induced consequences (See Fig. 3). The accumulation of BQA was most prominent, as it was consistently detected in the AKU model, whereas it was largely absent in the control condition. Low-level BQA production was observed in wt HepG2 cells only following supplementation with 4-HPP, likely due to saturation of endogenous HGD activity. Interestingly, BQA was still being produced in the AKU model even after deprivation of phenylalanine and tyrosine. Tyramine-O-sulfate was elevated upon tyrosine supplementation, yet it was completely abolished when cultured in phenylalanine/tyrosine-free medium. In contrast, metabolites associated with mitochondrial function and redox homeostasis, including L-acetylcarnitine, hydroxyisovaleric acid, and N-acetyl-serine, remained largely unaffected across the various culture conditions, indicating that these metabolic perturbations are largely independent of substrate availability and thus are the consequence of HGD deficiency.

**Fig. 3:**
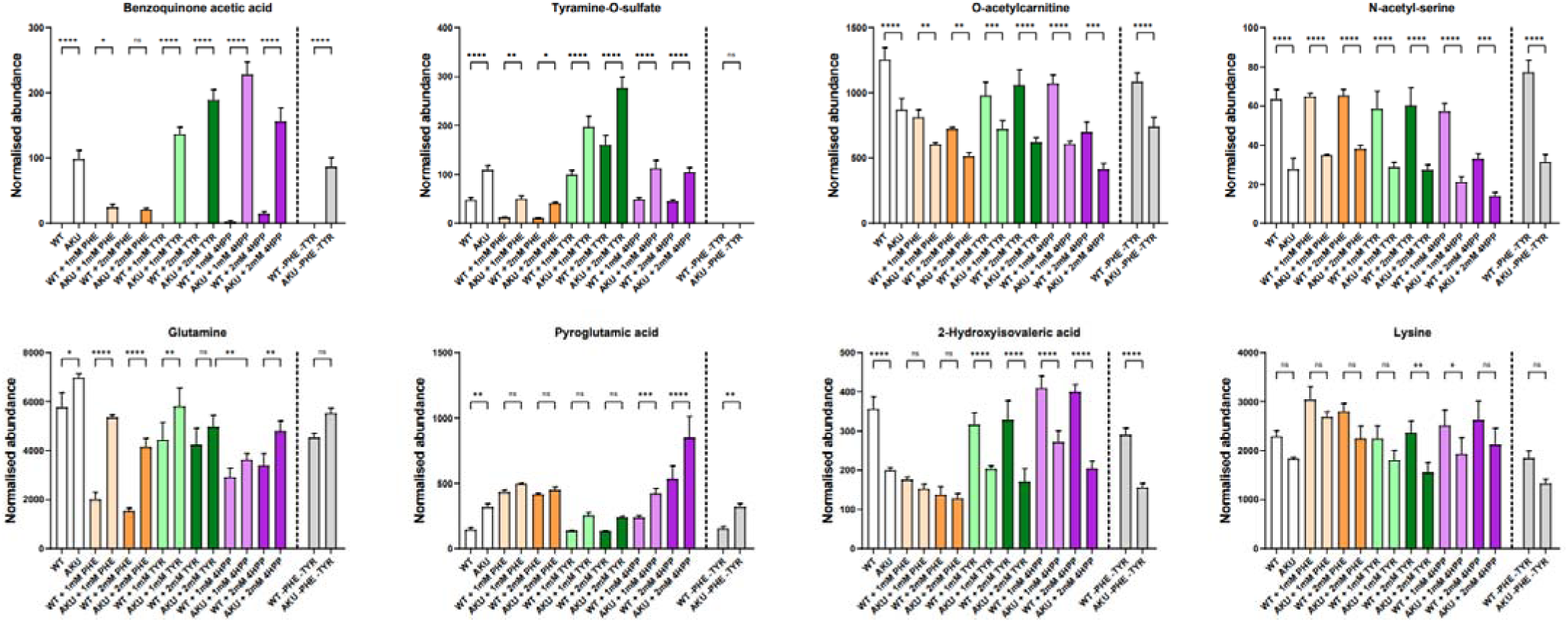
Metabolic profiling of wild type and HGD-deficient HepG2 cells under different cultivation conditions. Detection of metabolites in basal cultivation medium, exposed to 1mM and 2mM of phenylalanine, tyrosine, or 4-hydroxyphenylpyruvic acid, and in phenylalanine/tyrosine-free cultivation medium. One-way ANOVA (Šidák’s multiple comparison test): ns: not significant,*:p< 0.05, **: p < 0.01, ***: p < 0.001, ****: p < 0.0001 (n=3).

## 5. Discussion

In this study, we describe the first HGD-deficient hepatocyte-derived cell line, modelling the primary physiological site of tyrosine degradation and HGA production rather than the downstream osteoarticular manifestations examined in previous AKU models. Our findings support the view of AKU not solely as a defect in tyrosine degradation, but as a multi-system metabolic disorder characterized by widespread oxidative and chronic metabolic stress.

To generate this model, two universal HDR selection cassettes were developed that can be made gene-specific via a simple PCR^19^. Selection cassette integration was targeted to exon 3 of HGD, introducing a premature stop codon, with subsequent Cre recombinase-mediated excision of the selection markers yielding a footprint-free HGD gene KO. Although HDR genome editing efficiency is low and random selection cassette integration can occur, this comes at a much lower cost per gene KO. Delivery via ribonucleoprotein complex was favoured since it provides transient nuclease expression with reduced cell toxicity and unintended off-target activity. Loss of HGD protein was confirmed in 80% of clones and correct integration was confirmed by Western Blot, amplicon PCR, Sanger sequencing, and whole-genome sequencing, with no random integrations or chromosomal abnormalities detected. One acquired heterozygous ADGRG2 variant was identified, but this gene is not implicated in hepatic metabolism or AKU pathophysiology and showed no transcriptional alteration.

HGA readily undergoes auto-oxidation, generating free-radical intermediates that ultimately contribute to ochronotic pigment formation^3,22^. Although whether ochronotic pigment arises directly from HGA or through its proposed oxidation product BQA remains unresolved, the putative identification of BQA in our model, together with its detection following 4-HPP supplementation in wt HepG2 cells, provides biological support for its potential involvement in HGA oxidation and subsequently ochronotic pigment formation.

Consistent with Schiavone *et al.*^11^, HGD-deficient hepatocytes displayed impaired mitochondrial respiration. This was supported by altered regulators of mitochondrial biogenesis and homeostasis, downregulation of mitochondrial folate metabolism (SHMT2, MTHFD2, ALDH1L2), and increased ROS. Although DELE1 expression should increase under mitochondrial stress, OMA1, required for DELE1 cleavage and ISR activation, was downregulated and predicted to be inhibited^23^. Metabolomic profiling further indicated mitochondrial dysfunction, with accumulation of TCA-cycle intermediates, including malic and isocitric acid, and reductions in L-acetylcarnitine and N-acetylated amino acids, consistent with disturbed acetyl-CoA handling and oxidative metabolism^24^. This was accompanied by a coordinated metabolic shift toward increased glucose dependency and a partial uncoupling of glycolysis from oxidative phosphorylation, resembling a pseudohypoxic state. Accordingly, VHL, a major negative regulator of HIF-1α stabilization^25^, was predicted to be inhibited, while multiple important HIF-1α targets – including EGLN3, LOX, VEGF, EPO, PGK1, CA9, SLC16A3, BNIP3L, PDK1, and P4HA2 – were upregulated. Increased hippuric acid, uric acid, kynurenic acid, and hydroxybutyric acid levels independently supported a stress-adaptive phenotype associated with metabolic acidosis^26^, mitochondrial dysfunction^27^, reduced in oxidative phosphorylation^28^, and mitochondrial overload^29^, respectively. Increased kynurenic acid and indole-derived metabolites additionally indicated broader rewiring of tryptophan metabolism, consistent with rerouting toward the indolepyruvate pathway reported in AKU patients^30^.

Although pathway analysis predicted inhibition of the Warburg effect, this likely reflected reduced expression of proliferation-associated genes (GLS, GOT1, IDH2, MYC, and SLC1A5) rather than suppression of glycolytic adaptation. Together with reduced oxygen consumption, protein synthesis signatures, and proliferation, these findings suggest that HGD-deficient hepatocytes adopt a chronic stress-adaptive glycolytic survival state rather than a classical proliferation-associated Warburg phenotype, consistent with attenuated proliferation rate of cells exposed to HGA^11,15^. Extensive changes in amino acid transport and metabolite abundance further indicate amino acid pool imbalance.

Glycogenic enzymes GYS1 and GBE1 were also upregulated, suggesting increased energy-buffering capacity, whereas pentose phosphate pathway flux remained unchanged. An increased glycolysis flux is accompanied by an increase in the reactive byproduct methylglyoxal (MG), while its glutathione-dependent detoxification is potentially compromised by reduced cystine uptake (SLC7A11) and transsulfuration (CTH). Consistent with impaired glutathione synthesis, methionine and glutamate were depleted, whereas pyroglutamic acid accumulated, a metabolite associated with high-anion-gap metabolic acidosis^31^. MG accumulation promotes advanced glycation end-products (AGE) formation, which contributes to oxidative stress, lithiasis, tissue stiffening, and inflammation through protein cross-linking of structural proteins like collagen^33,34^, features also associated with AKU^4^. This is consistent with previous observations of reduced free thiols and increased protein carbonylation in *in vitro* AKU research^12,13^. Increased allysine, decreased lysine, and strong upregulation of LOX (converts lysine to allysine) further suggested activation of collagen and elastin cross-linking^35^, potentially representing an early metabolic signature of ochronotic extracellular-matrix remodelling.

Enhanced glycolysis may also increase lactate, which not only functions as a metabolic substrate but also as a regulator of chronic stress and cell-survival pathways^36^. Recently, lactate has been shown to modify histones through lysine lactylation, a dynamic post-translational modification associated with ferroptosis, an iron-dependent form of programmed cell death driven by lipid peroxidation^37^. In line with other research^8,12,13^, our HGD-deficient hepatic model also showed increased levels of lipid peroxidation alongside increased transferrin and transferrin receptor expression and marked SLC7A11 downregulation, all key features of ferroptosis. Although, the transcriptional expression of GPX4 remained unchanged. On a side note, lactylation at histone H4 K8 is tightly coupled with glycolysis, and more specifically PFKFB3^37^. Notably, AGE formation^38^, ferroptosis^39^, and lactylation^40^ have also been implicated in osteoarthritis, suggesting potential mechanistic links between hepatic metabolic dysfunction and downstream AKU pathology.

Canonical pathway and upstream-regulator analyses predicted inhibition of the integrated stress response (ISR), which normally acts to restore cellular homeostasis^41^. This may indicate that HGD-deficient cells undergo stable metabolic adaptation that reduces the requirement for sustained ISR activation. Despite extensive metabolic and extracellular-matrix remodelling, the hepatic model showed no clear inflammatory response, in contrast to the inflammation commonly observed in AKU patients^7,8^. Prolonged cultivation under constant tyrosine supplementation may have contributed to this adapted phenotype.

Finally, our data distinguish metabolites reflecting acute tyrosine pathway flux from those associated with the stable HGD-deficient state. BQA and tyramine-O-sulfate closely tracked substrate availability, whereas metabolites associated with mitochondrial dysfunction and oxidative stress remained altered irrespective of tyrosine flux. Thus, HGD deficiency establishes a persistent metabolic state extending beyond the immediate consequences of HGA accumulation. This distinction may also be relevant for biomarker development: flux-dependent metabolites may serve as diet-sensitive markers of tyrosine degradation, whereas consistently dysregulated metabolites may provide more robust disease-state biomarkers for monitoring AKU.

## 6. Conclusion

The present integrative multi-omic analysis demonstrates that HGD deficiency in a hepatocyte-derived cell line induces a broad chronic stress-adaptive metabolic program, extending far beyond impaired tyrosine catabolism. Collectively, the transcriptional and metabolomic signatures reveal coordinated mitochondrial dysfunction, pseudohypoxic glycolytic rewiring, disrupted amino acid homeostasis, impaired redox buffering capacity, and extensive extracellular matrix remodelling. Rather than supporting cellular growth, this metabolic state resembles a survival-oriented maintenance program characterized by reduced biosynthetic activity, altered nutrient transport, and functional rerouting of pyruvate metabolism away from mitochondrial oxidation toward oxygen-independent ATP production. Importantly, our model captures the metabolic consequences of endogenous HGD deficiency under chronic physiological tyrosine conditions, distinguishing it from other studies based primarily on exogenous HGA exposure. Additionally, this is the first HGD-deficient hepatocyte-derived cell line, thereby reflecting the primary physiological site of tyrosine degradation and HGA production rather than downstream osteoarticular manifestations commonly investigated in previous AKU models. These findings support the concept that AKU is not solely a defect of tyrosine degradation, but a multi-system metabolic disorder with widespread biochemical consequences due to oxidative and metabolic stress. Altogether, this study provides human-relevant insights into AKU pathophysiology and establishes a versatile platform for mechanistic investigation and therapeutic discovery.

## Manuscript details

1 Table & 2 Supplementary Tables

1 Supplementary Document

## Acknowledgements

This research was funded by the Research Foundation – Flanders (FWO) grant numbers 1S73019N, G023320N (EvolvAKUre), and G041521N (TALENT), Wetenschappelijk Fonds Willy Gepts (WFWG) from the UZ Brussel, VUB OZR grant, FWO medium-scale infrastructure funding (I001420N), and the Research Chair Mireille Aerens for Alternatives to Animal Testing.

## Declaration of generative AI and AI-assisted technologies in the writing process

During the preparation of this work, the authors used ChatGPT (GPT-5.5-mini, OpenAI) solely in order to improve the grammar, clarity, and readability of the manuscript. After using this tool, the authors reviewed and edited the content as needed and take full responsibility for the content of the published article.

## Author contributions statement

Conceptualization: M.R., J.H., G.B.G., J.D.K.

Investigation: M.R., G.D., B.N., A.H., G.A.

Methodology: M.R., G.D., S.L., B.N., J.N., G.A., J.D.K.

Resources: A.M., T.V., J.D.K.

Writing – Original Draft Preparation: M.R., J.D.K.

Writing – Review and Editing: M.R., G.D., S.L., B.N., L.D., N.S., J.H., G.A., T.V., G.B.G., J.D.K.

Visualization: M.R., J.D.K.

Supervision: J.D.K.

All authors have read and approved the final manuscript.

## Data Availability Statement

The Microarray data can be found on ArrayExpress via accession number E-MTAB-17171.

---

**Suppl. Fig. 1:**
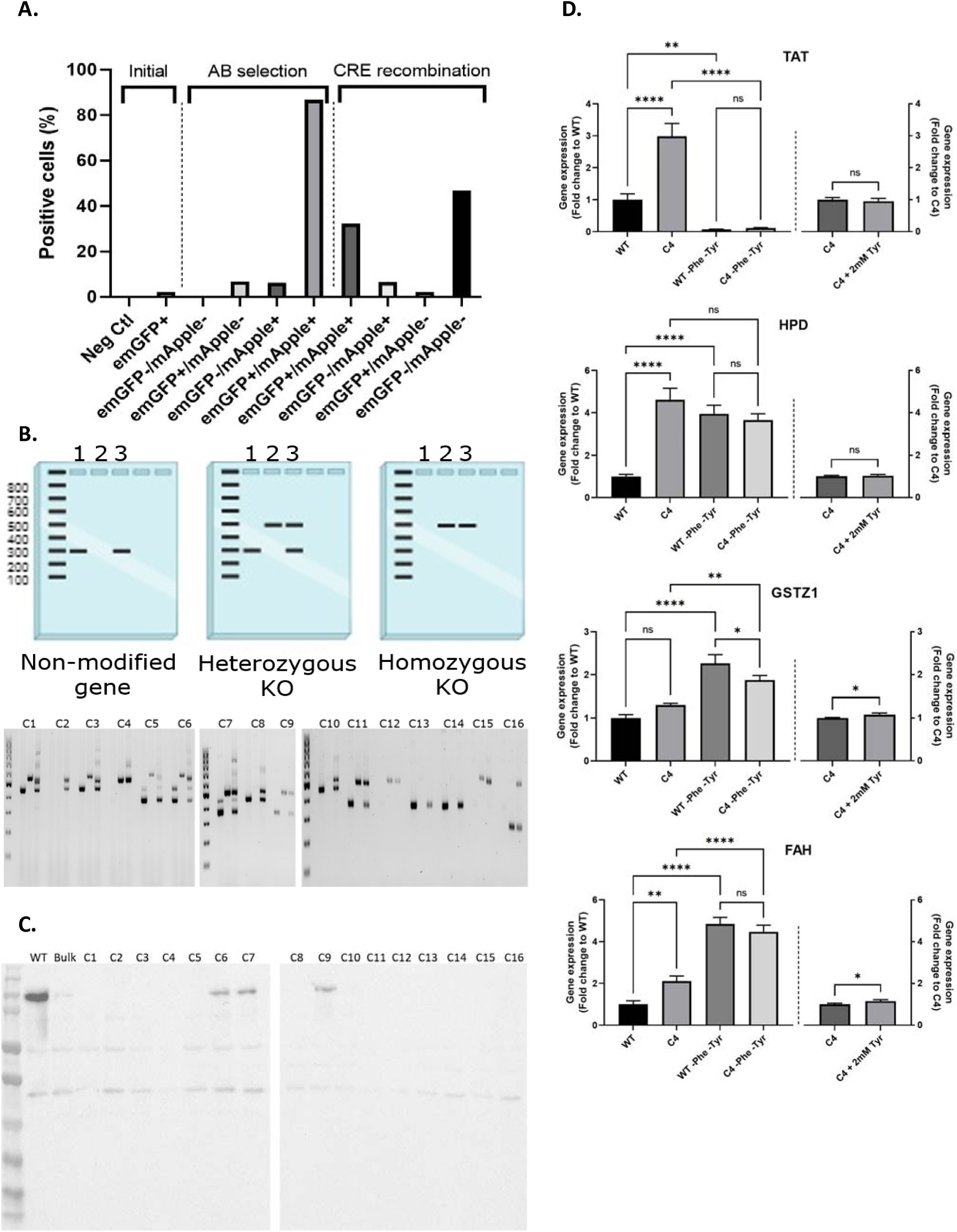
Generation and characterization of HGD-deficient HepG2 cells. **A:** Flow cytometric analysis of initial HDR efficiency before and after dual antibiotic selection, followed by Cre-recombinase induced excision of reporter cassette. **B:** Amplicon PCR method to verify the HGD knockout (KO) degree for sixteen indepedent clones: lane 1 = Fw+Rv primer, lane 2 = Fw+Jc primer, lane 3 = Fw+Rv+Jc primer. A non-modified gene results in one band at +/-300bp for lane 1 and 3, a heterozygous KO results in one band at +/-300bp for lane 1, one band at +/-500bp for lane 2, and two bands for lane 3 and a homozygous KO results in one at +/-500bp for lane 2 and 3. **C:** Full-length blot for HGD in HepG2 cells: Wild-type, bulk, and sixteen HGD KO clones are consecutively blotted. **D:** Relative TAT, HPD, GSTZ1, and FAH mRNA expression in HepG2 wt and HGD KO clone 4 measured by RT-qPCR and normalized to housekeeping genes GUSB, HPRT1, and UBC under supraphysiological levels of tyrosine and deprived from tyrosine and phenylalanine. One-way ANOVA (Tukey’s multiple comparison test) and Student’s t-test: ns: not significant, *:p <0.05, **: p < 0.01, ***: p < 0.001, ****: p < 0.0001 (n=3).

**Suppl. Fig. 2:**
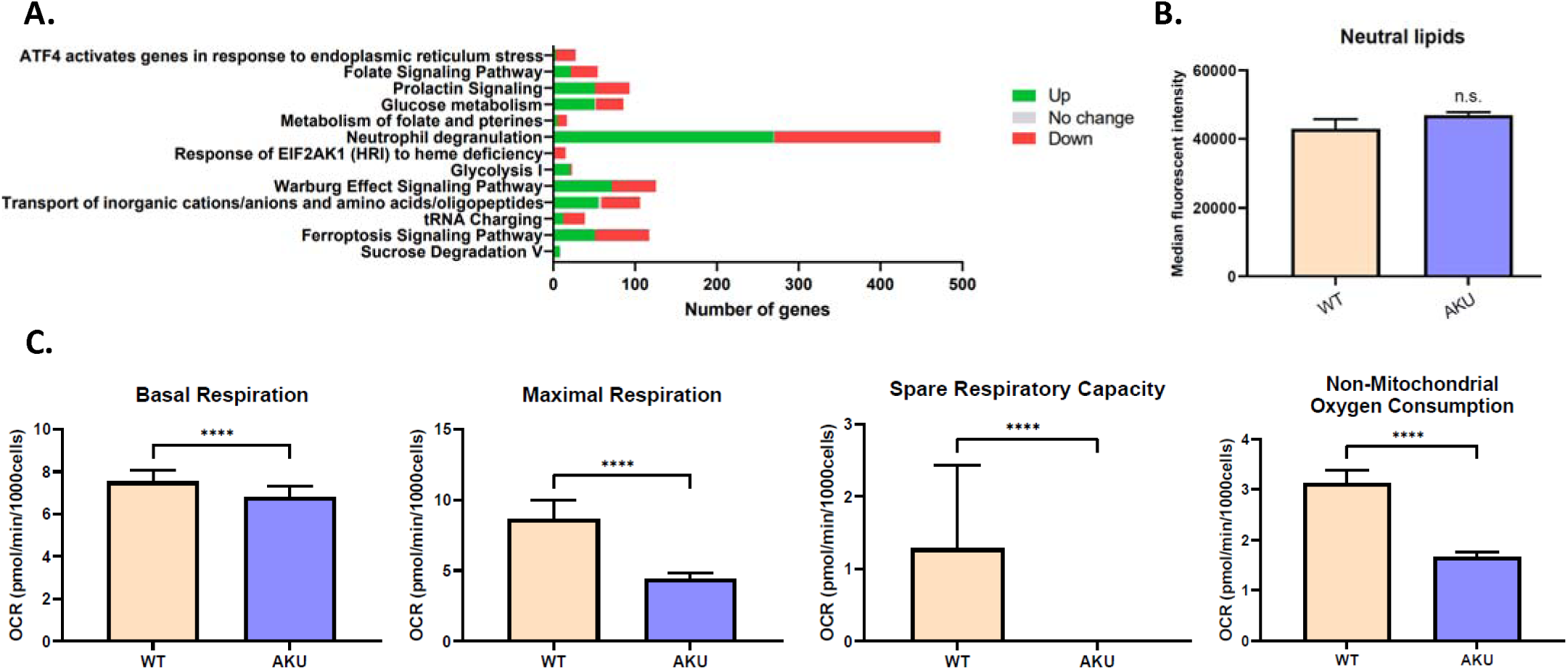
Characterization of wild type *versus* HGD-deficient HepG2 cells. **A:** Directionality and degree of modulation of canonical pathways due to HGD deficiency. **B:** Flow cytometric analysis of intracellular neutral lipids using Bodipy 493/503. student’s t-test: ns: not significant (n=3). **C:** Seahorse mitochondrial stress test: basal respiration, maximal respiration, spare respiratory capacity, and non-mitochondrial oxygen consumption of HepG2 cells versus HepG2 AKU model. Student’s t-test:****: p < 0.0001 (n=21).

## Bibliography

1. Zatkova, A., Ranganath, L. & Kadasi, L. Alkaptonuria: Current perspectives. Application of Clinical Genetics 13, (2020).

2. Ranganath, L. R., Norman, B. P. & Gallagher, J. A. Ochronotic pigmentation is caused by homogentisic acid and is the key event in alkaptonuria leading to the destructive consequences of the disease—A review. J. Inherit. Metab. Dis. 42, 776–792 (2019).

3. Grasso, D. et al. Ochronotic Deposition in Alkaptonuria: Semiquinone-Mediated Oxidative Coupling and Metabolic Drivers of Homogentisic Acid Accumulation. Int. J. Mol. Sci. 2025, *Vol.* 26, 26, 9674 (2025).

4. Bernardini, G. et al. Alkaptonuria. Nat. Rev. Dis. Prim. 10, 1–15 (2024).

5. Milella, M. S. et al. Alkaptonuria: From Molecular Insights to a Dedicated Digital Platform. Cells 13, 1–17 (2024).

6. Phornphutkul, C. et al. Natural History of Alkaptonuria. N. Engl. J. Med. 347, 2111–2121 (2002).

7. Spreafico, A. et al. Antioxidants inhibit SAA formation and pro-inflammatory cytokine release in a human cell model of alkaptonuria. Rheumatology (Oxford). 52, 1667 (2013).

8. Mastroeni, P. et al. An in vitro cell model for exploring inflammatory and amyloidogenic events in alkaptonuria. J. Cell. Physiol. 239, e31449 (2024).

9. Braconi, D. et al. Proteomic and redox-proteomic evaluation of homogentisic acid and ascorbic acid effects on human articular chondrocytes. J. Cell. Biochem. 111, 922–932 (2010).

10. Galderisi, S. et al. Homogentisic acid induces cytoskeleton and extracellular matrix alteration in alkaptonuric cartilage. J. Cell. Physiol. 236, 6011–6024 (2021).

11. Schiavone, M. L. et al. Mechanisms involved in the unbalanced redox homeostasis in osteoblastic cellular model of Alkaptonuria. Arch. Biochem. Biophys. 690, 108416 (2020).

12. Schiavone, M. L. et al. Homogentisic acid affects human osteoblastic functionality by oxidative stress and alteration of the Wnt/β-catenin signaling pathway. J. Cell. Physiol. 235, 6808–6816 (2020).

13. Braconi, D. et al. Redox-proteomics of the effects of homogentisic acid in an in vitro human serum model of alkaptonuric ochronosis. J. Inherit. Metab. Dis. 2011 346 34, 1163–1176 (2011).

14. Angeles, A. P., Badger, R., Gruber, H. E. & Seegmiller, J. E. Chondrocyte growth inhibition induced by homogentisic acid and its partial prevention with ascorbic acid. J. Rheumatol. 16, 512–517 (1989).

15. Tinti, L. et al. Development of an in vitro model to investigate joint ochronosis in alkaptonuria. Rheumatology 50, 271–277 (2011).

16. Norman, B. P. et al. Metabolomic studies in the inborn error of metabolism alkaptonuria reveal new biotransformations in tyrosine metabolism. Genes Dis. 9, 1129–1142 (2022).

17. Grasso, D., et al. Untargeted NMR Metabolomics Reveals Alternative Biomarkers and Pathways in Alkaptonuria. Int. J. Mol. Sci. 23, 15805 (2022).

18. Serafimov, K., Tischlarik, J. R. & Lämmerhofer, M. Targeted and untargeted urinary metabolomics of alkaptonuria patients using ultra high-performance liquid chromatography-tandem mass spectrometry. J. Pharm. Biomed. Anal. 256, 116684 (2025).

19. Liang, X., Potter, J., Kumar, S., Ravinder, N. & Chesnut, J. D. Enhanced CRISPR/Cas9-mediated precise genome editing by improved design and delivery of gRNA, Cas9 nuclease, and donor DNA. J. Biotechnol. 241, 136–146 (2017).

20. Ritz, C., Baty, F., Streibig, J. C. & Gerhard, D. Dose-Response Analysis Using R. PLoS One 10, e0146021 (2015).

21. Boeckmans, J. et al. Flow cytometric quantification of neutral lipids in a human skin stem cell-derived model of NASH. MethodsX 7, 101068 (2020).

22. Chow, W. Y., et al. Pigmentation Chemistry and Radical-Based Collagen Degradation in Alkaptonuria and Osteoarthritic Cartilage. Angew. Chemie -Int. Ed. 59, 11937–11942 (2020).

23. Yang, R. et al. De novo design of protein binders that target DELE1 to inhibit the mitochondrial stress response. bioRxiv Prepr. Serv. Biol. (2026).

24. McCann, M. R., De la Rosa, M. V. G., Rosania, G. R. & Stringer, K. A. L-Carnitine and Acylcarnitines: Mitochondrial Biomarkers for Precision Medicine. Metabolites 11, 51 (2021).

25. Basheeruddin, M. & Qausain, S. Hypoxia-Inducible Factor 1-Alpha (HIF-1α): An Essential Regulator in Cellular Metabolic Control. Cureus 16, e63852 (2024).

26. Shi, C., Guo, H. & Liu, X. High uric acid induced hippocampal mitochondrial dysfunction and cognitive impairment involving intramitochondrial NF-κB inhibitor α/nuclear factor-κB pathway. Neuroreport 33, 109–115 (2022).

27. Dzúrik, R., Spustová, V., Krivošíková, Z. & Gazíiková, K. Hippurate participates in the correction of metabolic acidosis. Kidney Int. 59, S278–S281 (2001).

28. Palzkill, V. R., Thome, T., Murillo, A. L., Khattri, R. B. & Ryan, T. E. Increasing plasma L-kynurenine impairs mitochondrial oxidative phosphorylation prior to the development of atrophy in murine skeletal muscle: A pilot study. Front. Physiol. 13, 992413 (2022).

29. Sousa, A. P. et al. Which Role Plays 2-Hydroxybutyric Acid on Insulin Resistance? Metabolites 11, 835 (2021).

30. Gertsman, I., Gangoiti, J. A., Nyhan, W. L. & Barshop, B. A. Perturbations of tyrosine metabolism promote the indolepyruvate pathway via tryptophan in host and microbiome. Mol. Genet. Metab. 114, 431–437 (2015).

31. Gamarra, Y. et al. Pyroglutamic acidosis by glutathione regeneration blockage in critical patients with septic shock. Crit. Care 23, 162 (2019).

32. Sbodio, J. I., Snyder, S. H. & Paul, B. D. Regulators of the transsulfuration pathway. Br. J. Pharmacol. 176, 583 (2018).

33. Prasad, C., Davis, K. E., Imrhan, V., Juma, S. & Vijayagopal, P. Advanced Glycation End Products and Risks for Chronic Diseases: Intervening Through Lifestyle Modification. Am. J. Lifestyle Med. 13, 384 (2017).

34. Vašková, J., Kováčová, G., Pudelský, J., Palenčár, D. & Mičková, H. Methylglyoxal Formation—Metabolic Routes and Consequences. Antioxidants 2025, Vol. 14, 14, 212 (2025).

35. Csiszar, K. Lysyl oxidases: A novel multifunctional amine oxidase family. Prog. Nucleic Acid Res. Mol. Biol. 70, 1–32 (2001).

36. Li, J., Ma, P., Liu, Z. & Xie, J. L- and D-Lactate: unveiling their hidden functions in disease and health. Cell Commun. Signal. 23, 134 (2025).

37. Lin, Z., Zou, Y., Zou, S. & Wen, K. Lactylation-mediated ferroptosis: A novel mechanism and therapeutic prospects in human diseases (Review). Int. J. Mol. Med. 57, 42 (2025).

38. He, C. P. et al. The role of AGEs in pathogenesis of cartilage destruction in osteoarthritis. Bone Jt. Res. 11, 292–300 (2022).

39. Liu, Y., Zhang, Z., Fang, Y., Liu, C. & Zhang, H. Ferroptosis in Osteoarthritis: Current Understanding. J. Inflamm. Res. 17, 8471 (2024).

40. Luan, S. & Luan, J. A multidimensional approach reveals the function of lactylation related genes in osteoarthritis. Sci. Reports 2025 151 15, 7743 (2025).

41. Pakos-Zebrucka, K., et al. The integrated stress response. EMBO Rep. 17, 1374 (2016).

