## Supplementary Document for "Unraveling the metabolic landscape of alkaptonuria through a human-relevant *in vitro* liver disease model"

Linker-LoxP-3xSTOP-EF1A-PuroR-P2A-emGFP-SV40-LoxP-3xSTOP-Uni

GGAAGTGGCTCAGGTTCTGGAATAACTTCGTATAGCATACATTATACGAAGTTATTAGATAACTGACGCGGCTCCGGTGCCCGTCAGTGGGCAGAGCGCACATCGCCCACAGTCCCCGAGAAGTTGGGGGGAGGGGTCGGCAATTGAACCGGTGCCTAGAGAAGGTGGCGCGGGGTAAACTGGGAAAGTGATGTCGTGTACTGGCTCCGCCTTTTTCCCGAGGGTGGGGGAGAACCGTATATAAGTGCAGTAGTCGCCGTGAACGTTCTTTTTCGCAACGGGTTTGCCGCCAGAACACAGGTAAGTGCCGTGTGTGGTTCCCGCGGGCCTGGCCTCTTTACGGGTTATGGCCCTTGCGTGCCTTGAATTACTTCCACCTGGCTGCAGTACGTGATTCTTGATCCCGAGCTTCGGGTTGGAAGTGGGTGGGAGAGTTCGAGGCCTTGCGCTTAAGGAGCCCCTTCGCCTCGTGCTTGAGTTGAGGCCTGGCCTGGGCGCTGGGGCCGCCGCGTGCGAATCTGGTGGCACCTTCGCGCCTGTCTCGCTGCTTTCGATAAGTCTCTAGCCATTTAAAATTTTTGATGACCTGCTGCGACGCTTTTTTTCTGGCAAGATAGTCTTGTAAATGCGGGCCAAGATCTGCACACTGGTATTTCGGTTTTTGGGGCCGCGGGCGGCGACGGGGCCCGTGCGTCCCAGCGCACATGTTCGGCGAGGCGGGGCCTGCGAGCGCGGCCACCGAGAATCGGACGGGGGTAGTCTCAAGCTGGCCGGCCTGCTCTGGTGCCTGGTCTCGCGCCGCCGTGTATCGCCCCGCCCTGGGCGGCAAGGCTGGCCCGGTCGGCACCAGTTGCGTGAGCGGAAAGATGGCCGCTTCCCGGCCCTGCTGCAGGGAGCTCAAAATGGAGGACGCGGCGCTCGGGAGAGCGGGCGGGTGAGTCACCCACACAAAGGAAAAGGGCCTTTCCGTCCTCAGCCGTCGCTTCATGTGACTCCACGGAGTACCGGGCGCCGTCCAGGCACCTCGATTAGTTCTCGAGCTTTTGGAGTACGTCGTCTTTAGGTTGGGGGGAGGGGTTTTATGCGATGGAGTTTCCCCACACTGAGTGGGTGGAGACTGAAGTTAGGCCAGCTTGGCACTTGATGTAATTCTCCTTGGAATTTGCCCTTTTTGAGTTTGGATCTTGGTTCATTCTCAAGCCTCAGACAGTGGTTCAAAGTTTTTTTCTTCCATTTCAGGTGTCGTGACAAGTTTGTACAAAAAAGCAGGCTGCCACCATGACCGAGTACAAGCCCACGGTGCGCCTCGCCACCCGCGACGACGTCCCCAGGGCCGTACGCACCCTCGCCGCCGCGTTCGCCGACTACCCCGCCACGCGCCACACCGTCGATCCGGACCGCCACATCGAGCGGGTCACCGAGCTGCAAGAACTCTTCCTCACGCGCGTCGGGCTCGACATCGGCAAGGTGTGGGTCGCGGACGACGGCGCCGCGGTGGCGGTCTGGACCACGCCGGAGAGCGTCGAAGCGGGGGCGGTGTTCGCCGAGATCGGCCCGCGCATGGCCGAGTTGAGCGGTTCCCGGCTGGCCGCGCAGCAACAGATGGAAGGCCTCCTGGCGCCGCACCGGCCCAAGGAGCCCGCGTGGTTCCTGGCCACCGTCGGCGTCTCGCCCGACCACCAGGGCAAGGGTCTGGGCAGCGCCGTCGTGCTCCCCGGAGTGGAGGCGGCCGAGCGCGCCGGGGTGCCCGCCTTCCTGGAGACCTCCGCGCCCCGCAACCTCCCCTTCTACGAGCGGCTCGGCTTCACCGTCACCGCCGACGTCGAGGTGCCCGAAGGACCGCGCACCTGGTGCATGACCCGCAAGCCCGGTGCCGGAAGCGGAGCCACGAACTTCTCTCTGTTAAAGCAAGCAGGAGATGTTGAAGAAAACCCCGGGCCTATGGTGAGCAAGGGCGAGGAGCTGTTCACCGGGGTGGTGCCCATCCTGGTCGAGCTGGACGGCGACGTAAACGGCCACAAGTTCAGCGTGTCCGGCGAGGGCGAGGGCGATGCCACCTACGGCAAGCTGACCCTGAAGTTCATCTGCACCACCGGCAAGCTGCCCGTGCCCTGGCCCACCCTCGTGACCACCTTCACCTACGGCGTGCAGTGCTTCGCCCGCTACCCCGACCACATGAAGCAGCACGACTTCTTCAAGTCCGCCATGCCCGAAGGCTACGTCCAGGAGCGCACCATCTTCTTCAAGGACGACGGCAACTACAAGACCCGCGCCGAGGTGAAGTTCGAGGGCGACACCCTGGTGAACCGCATCGAGCTGAAGGGCATCGACTTCAAGGAGGACGGCAACATCCTGGGGCACAAGCTGGAGTACAACTACAACAGCCACAAGGTCTATATCACCGCCGACAAGCAGAAGAACGGCATCAAGGTGAACTTCAAGACCCGCCACAACATCGAGGACGGCAGCGTGCAGCTCGCCGACCACTACCAGCAGAACACCCCCATCGGCGACGGCCCCGTGCTGCTGCCCGACAACCACTACCTGAGCACCCAGTCCGCCCTGAGCAAAGACCCCAACGAGAAGCGCGATCACATGGTCCTGCTGGAGTTCGTGACCGCCGCCGGGATCACTCTCGGCATGGACGAGCTGTACAAGTAAACCCAGCTTTCTTGTACAAAGTGGTGATGGCCGGCCGCTTCGAGCAGACATGATAAGATACATTGATGAGTTTGGACAAACCACAACTAGAATGCAGTGAAAAAAATGCTTTATTTGTGAAATTTGTGATGCTATTGCTTTATTTGTAACCATTATAAGCTGCAATAAACAAGTTAACAACAACAATTGCATTCATTTTATGTTTCAGGTTCAGGGGGAGGTGTGGGAGGTTTTTTAAAGCAAGTAAAACCTCTACAAATGTGGTAATAACTTCGTATAGCATACATTATACGAAGTTATTAGATAACTGACTCTGTATGCGATCGGCCAAG

Linker-LoxP-3xSTOP-EF1A-BsdR-P2A-mApple-SV40-LoxP-3xSTOP-Uni

GGAAGTGGCTCAGGTTCTGGAATAACTTCGTATAGCATACATTATACGAAGTTATTAGATAACTGACGCGGCTCCGGTGCCCGTCAGTGGGCAGAGCGCACATCGCCCACAGTCCCCGAGAAGTTGGGGGGAGGGGTCGGCAATTGAACCGGTGCCTAGAGAAGGTGGCGCGGGGTAAACTGGGAAAGTGATGTCGTGTACTGGCTCCGCCTTTTTCCCGAGGGTGGGGGAGAACCGTATATAAGTGCAGTAGTCGCCGTGAACGTTCTTTTTCGCAACGGGTTTGCCGCCAGAACACAGGTAAGTGCCGTGTGTGGTTCCCGCGGGCCTGGCCTCTTTACGGGTTATGGCCCTTGCGTGCCTTGAATTACTTCCACCTGGCTGCAGTACGTGATTCTTGATCCCGAGCTTCGGGTTGGAAGTGGGTGGGAGAGTTCGAGGCCTTGCGCTTAAGGAGCCCCTTCGCCTCGTGCTTGAGTTGAGGCCTGGCCTGGGCGCTGGGGCCGCCGCGTGCGAATCTGGTGGCACCTTCGCGCCTGTCTCGCTGCTTTCGATAAGTCTCTAGCCATTTAAAATTTTTGATGACCTGCTGCGACGCTTTTTTTCTGGCAAGATAGTCTTGTAAATGCGGGCCAAGATCTGCACACTGGTATTTCGGTTTTTGGGGCCGCGGGCGGCGACGGGGCCCGTGCGTCCCAGCGCACATGTTCGGCGAGGCGGGGCCTGCGAGCGCGGCCACCGAGAATCGGACGGGGGTAGTCTCAAGCTGGCCGGCCTGCTCTGGTGCCTGGTCTCGCGCCGCCGTGTATCGCCCCGCCCTGGGCGGCAAGGCTGGCCCGGTCGGCACCAGTTGCGTGAGCGGAAAGATGGCCGCTTCCCGGCCCTGCTGCAGGGAGCTCAAAATGGAGGACGCGGCGCTCGGGAGAGCGGGCGGGTGAGTCACCCACACAAAGGAAAAGGGCCTTTCCGTCCTCAGCCGTCGCTTCATGTGACTCCACGGAGTACCGGGCGCCGTCCAGGCACCTCGATTAGTTCTCGAGCTTTTGGAGTACGTCGTCTTTAGGTTGGGGGGAGGGGTTTTATGCGATGGAGTTTCCCCACACTGAGTGGGTGGAGACTGAAGTTAGGCCAGCTTGGCACTTGATGTAATTCTCCTTGGAATTTGCCCTTTTTGAGTTTGGATCTTGGTTCATTCTCAAGCCTCAGACAGTGGTTCAAAGTTTTTTTCTTCCATTTCAGGTGTCGTGACAAGTTTGTACAAAAAAGCAGGCTGCCACCATGGCCAAGCCTTTGTCTCAAGAAGAATCCACCCTCATTGAAAGAGCAACGGCTACAATCAACAGCATCCCCATCTCTGAAGACTACAGCGTCGCCAGCGCAGCTCTCTCTAGCGACGGCCGCATCTTCACTGGTGTCAATGTATATCATTTTACTGGGGGACCTTGTGCAGAACTCGTGGTGCTGGGCACTGCTGCTGCTGCGGCAGCTGGCAACCTGACTTGTATCGTCGCGATCGGAAATGAGAACAGGGGCATCTTGAGCCCCTGCGGACGGTGCCGACAGGTGCTTCTCGATCTGCATCCTGGGATCAAAGCCATAGTGAAGGACAGTGATGGACAGCCGACGGCAGTTGGGATTCGTGAATTGCTGCCCTCTGGTTATGTGTGGGAGGGCGGAAGCGGAGCCACGAACTTCTCTCTGTTAAAGCAAGCAGGAGATGTTGAAGAAAACCCCGGGCCTATGGTGAGCAAGGGCGAGGAGAATAACATGGCCATCATCAAGGAGTTCATGCGCTTCAAGGTGCACATGGAGGGCTCCGTGAACGGCCACGAGTTCGAGATCGAGGGCGAGGGCGAGGGCCGCCCCTACGAGGCCTTTCAGACCGCTAAGCTGAAGGTGACCAAGGGTGGCCCCCTGCCCTTCGCCTGGGACATCCTGTCCCCTCAGTTCATGTACGGCTCCAAGGTCTACATTAAGCACCCAGCCGACATCCCCGACTACTTCAAGCTGTCCTTCCCCGAGGGCTTCAGGTGGGAGCGCGTGATGAACTTCGAGGACGGCGGCATTATTCACGTTAACCAGGACTCCTCCCTGCAGGACGGCGTGTTCATCTACAAGGTGAAGCTGCGCGGCACCAACTTCCCCTCCGACGGCCCCGTAATGCAGAAGAAGACCATGGGCTGGGAGGCCTCCGAGGAGCGGATGTACCCCGAGGACGGCGCCCTGAAGAGCGAGATCAAGAAGAGGCTGAAGCTGAAGGACGGCGGCCACTACGCCGCCGAGGTCAAGACCACCTACAAGGCCAAGAAGCCCGTGCAGCTGCCCGGCGCCTACATCGTCGACATCAAGTTGGACATCGTGTCCCACAACGAGGACTACACCATCGTGGAACAGTACGAACGCGCCGAGGGCCGCCACTCCACCGGCGGCATGGACGAGCTGTACAAGTAAACCCAGCTTTCTTGTACAAAGTGGTGATGGCCGGCCGCTTCGAGCAGACATGATAAGATACATTGATGAGTTTGGACAAACCACAACTAGAATGCAGTGAAAAAAATGCTTTATTTGTGAAATTTGTGATGCTATTGCTTTATTTGTAACCATTATAAGCTGCAATAAACAAGTTAACAACAACAATTGCATTCATTTTATGTTTCAGGTTCAGGGGGAGGTGTGGGAGGTTTTTTAAAGCAAGTAAAACCTCTACAAATGTGGTAATAACTTCGTATAGCATACATTATACGAAGTTATTAGATAACTGACTCTGTATGCGATCGGCCAAG
