## Supplementary Table 1 for "Unraveling the metabolic landscape of alkaptonuria through a human-relevant *in vitro* liver disease model"

**Suppl. Table 1: Comparative transcriptomic analysis of HGD-deficient HepG2 cells (AKU model) versus wild type HepG2 cells.** Showing differentially expressed genes (DEGs) with an absolute fold change >10. Yellow background indicates increased gene expression, whereas blue background indicates decreased gene expression.

| **Gene** | **Gene name** | **Function** | **Fold change** |
| --- | --- | --- | --- |
| RAP1A | RAP1A, Member Of RAS Oncogene Family | Cell signaling | 308.88 |
| TNNI2 | Troponin I2, Fast Skeletal Type | Muscle contraction regulation | 34.94 |
| FAM13A | Family With Sequence Similarity 13 Member A | Cell signaling | 32.43 |
| SLC2A3 | Solute Carrier Family 2 Member 3 | Glucose transporter | 25.76 |
| CA9 | Carbonic Anhydrase 9 | pH regulation | 25.57 |
| TUBB1 | Tubulin Beta 1 Class VI | Cytoskeletal organization | 25.17 |
| EGLN3 | Egl-9 Family Hypoxia Inducible Factor 3 | HIF signaling regulation | 19.06 |
| ADSS1 | Adenylosuccinate Synthase 1 | Purine nucleotide biosynthesis | 16.08 |
| SLC2A14 | Solute Carrier Family 2 Member 14 | Glucose transporter | 15.49 |
| SLC51A | Solute Carrier Family 51 Member A | Bile transporter | 14.81 |
| SLC6A8 | Solute Carrier Family 6 Member 8 | Creatine transporter | 13.60 |
| LCN15 | Lipocalin 15 | Small molecule transport | 13.33 |
| CHST15 | Carbohydrate Sulfotransferase 15 | ECM remodeling | 12.39 |
| NEGR1 | Neuronal Growth Regulator 1 | Cell adhesion | 12.36 |
| NDRG1 | N-Myc Downstream Regulated 1 | Stress response | 11.65 |
| PFKFB3 | 6-Phosphofructo-2-Kinase/Fructose-2,6-Biphosphatase 3 | Glycolysis regulation | 11.35 |
| TMEM37 | Transmembrane Protein 37 | Calcium transporter | 11.35 |
| LOX | Lysyl Oxidase | ECM remodeling | 11.00 |
| SPAG4 | Sperm Associated Antigen 4 | Cytoskeletal organization | 10.42 |
| SLC7A10 | Solute Carrier Family 7 Member 10 | Neutral amino acid transporter | 10.10 |
| ALDH1L2 | Aldehyde Dehydrogenase 1 Family Member L2 | Mitochondrial folate metabolism | -10.09 |
| SPP1 | Secreted Phosphoprotein 1 | Cell adhesion | -10.39 |
| AREG | Amphiregulin | Cell proliferation and survival | -11.78 |
| CTH | Cystathionine Gamma-Lyase | Cysteine production | -14.21 |
| ASNS | Asparagine Synthetase (Glutamine-Hydrolyzing) | Asparagine biosynthesis | -15.04 |
| HGD | Homogentisate 1,2-Dioxygenase | Tyrosine degradation | -17.15 |
| SLC7A11 | Solute Carrier Family 7 Member 11 | Cystine/glutamate antiporter | -17.59 |
