## Supplementary Table 2 for "Unraveling the metabolic landscape of alkaptonuria through a human-relevant *in vitro* liver disease model"

**Suppl. Table 2: Upstream regulator analysis of HGD-deficient HepG2 cells (AKU model) versus wild-type HepG2 cells.** Showing upstream regulators with a q-value <0.05 and a predicted activation state. Yellow background indicates a predicted activated state, whereas blue background indicates a predicted inhibited state.

| **Upstream  Regulator** | **Fold  Change** | **Molecule Type** | **Predicted  Activation  State** | **Activation  z-score** | **p-value of  overlap** | **Target Molecules** |
| --- | --- | --- | --- | --- | --- | --- |
| TFAM | -2.42 | transcription regulator | Activated | 3.74 | 1.77E-12 | ALDH1L2, ASNS, CARS1, CTH, GARS1, MARS1, MTHFD2, SARS1, SHMT2, SLC3A2, SLC51A, SLC7A1, SLC7A11, YARS1 |
| HELLS | -1.49 | enzyme | Activated | 3.16 | 3.39E-11 | ASNS, CARS1, CTH, GARS1, MARS1, MTHFD2, SARS1, SHMT2, SLC3A2, YARS1 |
| MIMS1 | 1.31 | other | Activated | 2.80 | 2.47E-07 | ASNS, CARS1, GARS1, MARS1, SARS1, SLC3A2, SLC7A11, YARS1 |
| LONP1 | 1.07 | peptidase | Activated | 2.45 | 2.19E-06 | ALDH1L2, ASNS, GARS1, MARS1, MTHFD2, SARS1, SOD2 |
| ARID1A | 1.50 | transcription regulator | Activated | 2.35 | 2.37E-05 | ABCG2, AGRN, CKB, CLGN, ID2, JUP, KDELR3, LOX, NREP, P4HA2, PCOLCE2, RAP1A, SLC2A3, SYNE2 |
| ALDH2 | -1.45 | enzyme | Activated | 2.24 | 5.88E-05 | CTH, MTHFD2, SHMT2, SLC3A2, SLC7A11 |
| TRAP1 | -1.17 | enzyme | Activated | 2.45 | 6.58E-05 | ALDH1L2, GARS1, MARS1, MTHFD2, SARS1, SLC2A3 |
| P2RX7 | -1.31 | ion channel | Activated | 2.21 | 7.50E-05 | ALDOA, ALDOC, IDH2, PFKL, TPI1 |
| DELE1 | 1.29 | other | Inhibited | -3.60 | 1.17E-16 | ALDH1L2, ASNS, CARS1, CTH, GARS1, MARS1, MTHFD2, SARS1, SHMT2, SLC3A2, SLC7A1, SLC7A11, YARS1 |
| ATF4 | -2.23 | transcription regulator | Inhibited | -3.75 | 2.66E-16 | ABCG2, ASNS, CARS1, CEBPB, CEBPG, CTH, FYN, GARS1, LGALS3, MAP1LC3B, MARS1, MTHFD2, SARS1, SHMT2, SLC1A5, SLC38A1, SLC3A2, SLC7A1, SLC7A11, SOD2 |
| OMA1 | -1.66 | peptidase | Inhibited | -2.80 | 3.63E-10 | ALDH1L2, ASNS, CEBPG, CTH, GARS1, MTHFD2, MYC, SHMT2 |
| KRAS | -1.98 | enzyme | Inhibited | -2.02 | 1.71E-09 | ADH4, AGRN, ALDOA, ALDOC, CEBPB, CTH, CYP24A1, DDR1, FKBP11, ITIH5, JUP, LGALS3, LOX, MAP1LC3B, MMP3, MYC, NR5A2, PALLD, PCOLCE2, PFKL, PHLDB2, PLAGL1, SCARB1, SLC3A2, SLC7A1, SLC7A11, STAT2, TCEA1, TPI1 |
| EIF2AK3 | 1.10 | kinase | Inhibited | -2.16 | 1.95E-08 | ASNS, CARS1, CEBPB, CEBPG, ERN1, MAP1LC3B, MYC, SARS1, SHMT2, SLC7A1, YARS1 |
| NTRK1 | 1.26 | kinase | Inhibited | -2.18 | 5.31E-08 | ALDH1L2, ASNS, CARS1, CEBPB, MTHFD2, MYO10, P4HA2, PHLDB2, SARS1, SHMT2, SLC3A2, SOD2, SYNE2, YARS1 |
| UCP1 | 1.13 | transporter | Inhibited | -3.094 | 6.78E-07 | ASNS, CEBPG, CTH, ERN1, GARS1, GOT1, MTHFD2, MYC, SHMT2, SLC3A2 |
| PGR | -1.47 | ligand-dependent nuclear receptor | Inhibited | -2.331 | 5.09E-05 | ABCG2, CEBPB, CLDN6, DDX21, GLS, GOT1, LOX, MPZL2, MYC, MYO10, P4HA2, RHOBTB3, TSC22D3 |
| IL1B | -1.29 | cytokine | Inhibited | -2.142 | 2.61E-04 | ABCG2, APOL1, CEBPB, CEBPG, CPB2, CYP24A1, EPO, ID2, IFRD1, JUP, KYNU, LOX, MMP3, MYC, P4HA2, RXRA, SCARB1, SLC11A2, SLC37A4, SLC7A11, SOD2, TSC22D3 |
| VHL | -1.07 | transcription regulator | Inhibited | -2.21 | 6.37E-04 | C5AR1, EPO, LOX, NREP, SLC11A2, SLC2A3, SOD2 |
| WBP2 | 1.24 | transcription regulator | Inhibited | -2.22 | 3.05E-03 | CTH, CYP24A1, JDP2, SFXN3, SLC7A11 |
| EGFR | 2.96 | kinase | Inhibited | -2.12 | 3.55E-03 | ABCG2, ACSL3, CEBPB, DDX21, FKBP11, IFRD1, MYC, SCARB1, SLC2A3, SLC37A4, SLC7A1, SLC7A11 |
| STAT3 | 1.45 | transcription regulator | Inhibited | -2.38 | 9.95E-03 | BEX2, C5AR1, CEBPB, ID2, IL21R, LOX, MMP3, MYC, PCOLCE2, PFKL, PLAGL1, SOD2, STAT2 |
| ERG | 1.25 | transcription regulator | Inhibited | -2.00 | 1.29E-02 | CEBPG, DIAPH2, FYN, MMP3, MPZL2, MYO10, PHLDB2, TSC22D3 |
| HRAS | 1.19 | enzyme | Inhibited | -2.20 | 3.23E-02 | ASNS, BOK, CEBPB, CYP24A1, ID2, JUP, MMP3, MYC, PLOD1, UBASH3B |
| RELA | 1.29 | transcription regulator | Inhibited | -2.57 | 3.78E-02 | ABCG2, BEX2, CEBPB, MMP3, MYC, SLC2A3, SOD2, TANK |
| TREM1 | 1.13 | Transmembrane receptor | Inhibited | -2.00 | 5.69E-02 | ACSL3, ASNS, CEBPB, RHOBTB3 |
